# The metabolic and molecular mechanisms underlying running-induced energy compensation

**DOI:** 10.1101/2025.06.09.658628

**Authors:** Paula Sanchis, Cecilie Bæch-Laursen, Joel Vizueta, Hazal Haytural, Nigel Kurgan, Kaspar W. Persson, Alexander G. Bjørnholt, Rodrigo Salinas, Diana Samodova-Sommer, Juen Guo, Thomas E. Jensen, Atul Deshmukh, Bente K. Pedersen, Claus Brandt

## Abstract

Exercise not only regulates energy expenditure but also appetite, yet the underpinnings remain unclear. We describe that increased energy intake is a defense against energy loss that depends on initial running distance and operates independently of diet and age. Running caused a rapid circadian-dependent fat remodeling leading to a decline in circulating leptin accompanied by the activation of hypothalamic neurons. We discovered that the activation of the β3-adrenergic receptor drives running-induced fat loss and the lower leptin triggers energy compensation by upregulating *Neuropeptide Y*. Once energy compensation is achieved, running is associated with molecular changes in hypothalamic signaling related to appetite and functional adaptations, such as enhanced sensitivity to hunger and satiety signals and increased responsiveness to appetite suppression induced by α-Melanocyte-Stimulating Hormone. The increased food intake persisted without fat rebound beyond running in both lean and obese young mice, uncovering a new homeostatic balance in young mice.

---

Energy compensation is the bodily response that works to minimize the energy gap of increased energy expenditure (*1*). A decrease in energy expended on other biological processes and an increase in caloric intake have been postulated as compensatory mechanisms of energy expended by exercise (*2–4*). Energy intake follows a J-shaped relationship with physical activity, where low activity is inversely correlated with intake, while moderate to high activity shows a positive association (*5*, *6*). Such responses to physical activity are highly disadvantageous when the goal of exercising is to decrease adiposity and body mass. Exercising also promotes subjective hunger, enhances post-meal satiety and reduces susceptibility to overconsumption, revealing that both hunger and satiety are better regulated in active individuals (*4*, *7*). Thus, in addition to expending calories, physical activity may increase feeding sensitivity by a better coupling of energy intake to energy expenditure (*8*, *9*).

Similar to humans, voluntary wheel running regulates food intake and prevents overconsumption of high-fat-diet (HFD) in rodents (*10–12*). In mice, feeding behavior follows a circadian pattern, higher during the active dark phase and minimal during the resting light phase. In chronic running mice, food intake increases, particularly before and during the latter part of the dark phase (*13*). Running also decreases body mass and adiposity and increases lean mass in a distance-, duration-, and intensity-dependent manner (*12*). Adipose tissue is positioned as the crucial peripheral organ that regulates body weight through its action on the hypothalamus, the central regulator of energy balance. In particular, leptin is an anorexigenic hormone produced mainly in adipose cells that binds to its receptor in the hypothalamus and influences food intake (*14*, *15*). In the arcuate nucleus of hypothalamus (ARC), circulating leptin activates anorexigenic pro-opiomelanocortin (POMC)/cocaine-amphetamine-regulated transcript (CART) neurons, leading to the secretion of α-Melanocyte-stimulating hormone (α-MSH) (*16*), which then inhibits food intake by binding to melanocortin 4 receptor in paraventricular hypothalamic nucleus (PVN) neurons. Moreover, leptin suppresses the activation of Agouti-related protein (AGRP)/ Neuropeptide Y (NPY) neurons, further contributing to the inhibition of food intake (*17–19*). Of note, exercise reduces the leptin levels, correlating with changes in total fat, and enhances central leptin signaling (*11*, *20–22*). Emerging data from human experiments reveal an increase in the circulating ratio of adipocyte-derived adiponectin and leptin following exercise (*23*). Adiponectin has been described as a starvation signal via AdipoR1 opposing the role of leptin in the hypothalamus,(*24*) although its role in exercised-induced energy compensation has not been explored. Beyond food intake, leptin modulates energy expenditure, glucose homeostasis and lipolysis. The latter is partially mediated by the β-adrenergic system (*25–27*). The stimulation of β3-adrenergic receptor (AR) suppresses *leptin* expression in white fat (*28*). Interestingly, β3-AR is recognized as a major receptor for norepinephrine in mouse adipocytes and its pharmacological blockage before acute exercise prevents *leptin* downregulation in rat retroperitoneal fat, but not in epididymal fat (*29*). However, the role of β3-AR in running-induced fat mass loss and energy compensation is unknown. Thus, despite decades of scrutiny of exercise-induced energy compensation, a detailed physiological and molecular understanding is still lacking.

Here, we reveal that energy compensation is a conserved response associated to increased energy expenditure from running and proceeded by temporal morphological and molecular circadian-effect changes in subcutaneous and visceral fat, reduced circulating leptin and simultaneous changes in chronic hypothalamic neuronal activity. The initial running distance drives β3-AR-mediated fat loss and the decline of circulating leptin triggers running-induced energy compensation by upregulating *NPY*. After energy compensation, increased food intake remained relatively constant with improved appetite regulation, changes in the hypothalamic hunger signaling and minimal additional fat loss during running. This adaptive response to increased food intake and lower fat mass was maintained during inactivity in lean and obese young mice, and it was associated with hypothalamic adaptations. These findings reveal the pivotal role of the adrenergic system and leptin in mediating running-induced energy compensation and underscore the new homeostatic steady state in young mice after running.

## RESULTS

### Running-induced fat loss drives increased food intake and subsequent gain in lean mass

We monitored single-housed *ad libitum*-chow-fed male mice assigned by baseline traits to active (functional wheel) or sedentary (blocked wheel) groups for 5 weeks (Fig. 1A-B). Running resulted in an initial body weight loss and linear loss of fat mass reaching -0.50±0.09 in the first 4 days, which was compensated by an increase in food intake preventing excessive further fat loss, which reached -0.62±0.1 (Fig. 1C-E). After the increase in food intake, changes in fat increased in parallel with those of sedentary mice due to somatic growth, with a slight potentiation in lean mass growth due to running (Fig. 1E-F).

**Fig. 1.**
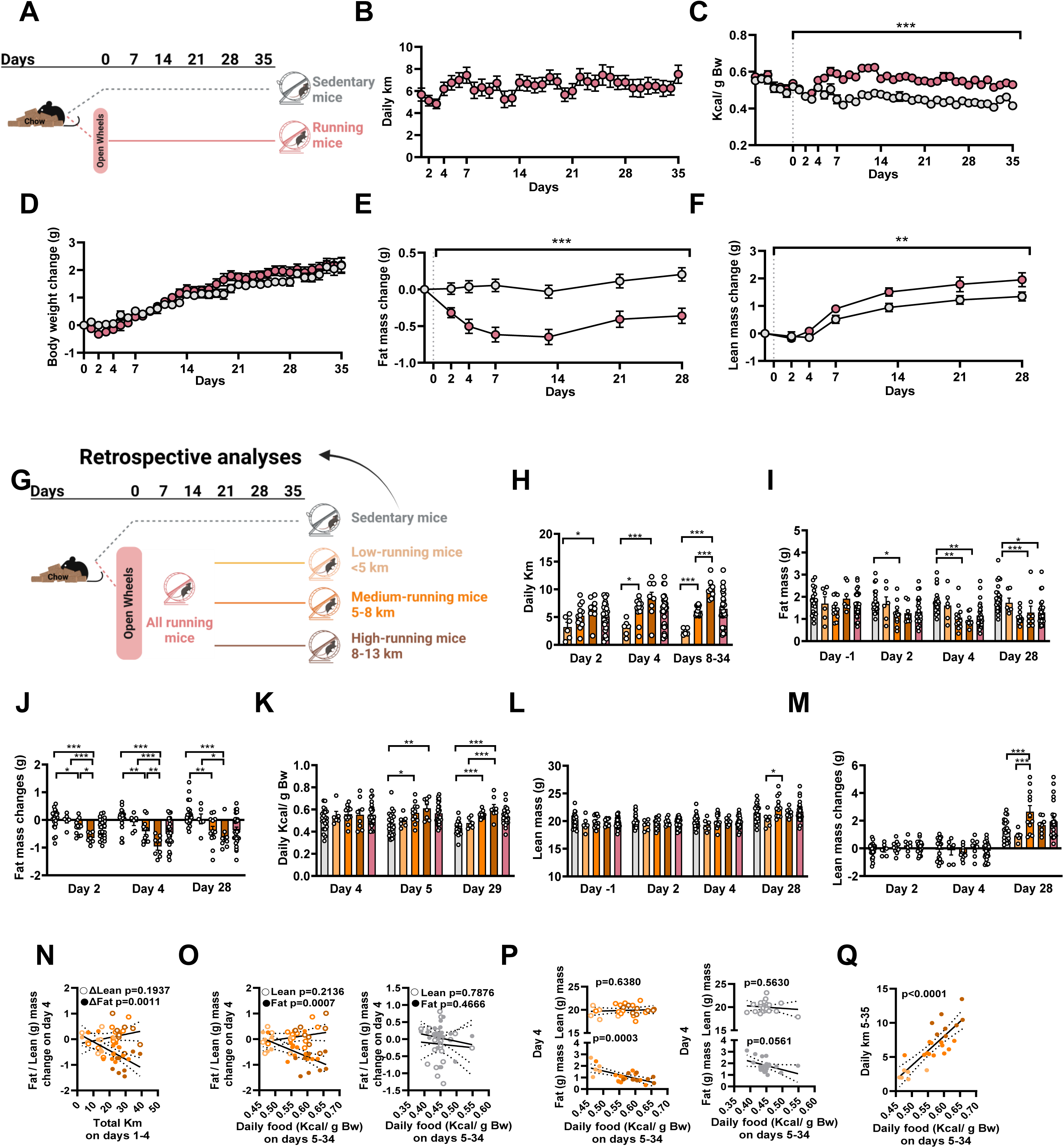
**Running yields fat loss and then increased food intake. A-F**) Chronic running. **A**, Schematic. **B**, Distance. **C**, Food intake. **D**, Body weight changes. **E**, Fat changes. **F**, Lean changes. **G-M**) Retrospective study. **G**, Schematic. **H**, Distance. **I**, Fat mass. **J**, Fat changes. **K**, Food intake. **L**, Lean mass. **M**, Lean changes. Pink bars were excluded from the analysis. **N-Q**) Associations. **N**, Distance for 4 days and fat/lean mass changes. **O**, Food intake and fat/lean changes in running (left) and sedentary mice (right) on day 4. **P**, Food intake and fat/lean in running (left) and sedentary mice (right) on day 4. **Q**, Daily intake and running. Data were analyzed using a one-way repeated-measures analysis of variance (ANOVA) to determine the main effects of running (*) in **C**-**F**. However, whether Mauchly’s sphericity test and/or Shapiro-Wilk’s Normality test was significant, then repeated-measures Generalized Estimating Equations (GEE) was used. One-way ANOVA with Bonferroni post hoc multiple comparison tests was conducted to evaluate the effect of running (*), but whether Shapiro-Wilk’s test and/or Levene’s test for homogeneity of variances were significant, we used Generalized Linear Model (GzLM) with sequential Bonferroni post hoc multiple comparisons (**H**-**M**). Simple linear regression was used in **N**-**Q.** Data are mean±SEM. *P≤0.05, **P≤0.01, ***P≤0.001. **A** and **G** were created using Biorender.

Running mice varied in distance covered. To address how distance, and indirectly energy expenditure, are related to energy compensation (Fig. 1G), we categorized them into three groups based on distance covered during a stable running period (8-35 days). Mice that ran 2.45±0.22 km/day, named low-runners, did not lose fat and thus did not compensate. In contrast, medium-runners, and high-runners, which ran 6.16±0.24km/day and 10.04±0.58km/day over a stable running period, respectively, exhibited a proportional loss in fat mass and increase in food intake (Fig. 1H-I). The medium-runners and high-runners lost 0.41±0.11g and 0.96±0.12g fat mass, respectively, on day 4 and ate 0.57±0.03 and 0.61±0.03 g/body weight, respectively, the following day (Fig. 1I-K). Minor differences were observed in lean mass (Fig. 1L-M), and the total running distance until day 4 was correlated with fat loss, but not with lean mass (Fig. 1N). Moreover, fat loss, but not lean gain, positively correlated with daily intake. This suggests a prominent role of fat loss in the response to running-induced food intake (Fig. 1O). Interestingly, fat mass in running mice on day 4 was negatively associated with daily food intake whereas no association was observed with lean mass. The same trend was observed in sedentary mice (Fig. 1P). From day 4 onward, the daily food intake positively correlated with the previous day’s running distance (Fig. S1A) resulting in a significant correlation between the average daily running distance and food intake across the experiment (Fig. 1Q). These findings highlight that the body senses and matches energy intake to energy expended by running, but only above a certain threshold.

### Energy compensation operates independently of diet, age and sub-strains in lean mice

The energy compensation was further explored in response to a calorie-dilution diet, which contains indigestible components,(*30*) to evaluate its conservation in lean mice with different energy stores. First, 6-week-old C57BL/6NRj mice were *ad libitum* fed with control-diet or calorie-dilution-diet for 2 weeks and then, blocked or functional wheels were provided for 5 weeks. Using a threshold of 5 km during the stable active period (days 8–35) as a trigger food compensation observed in our lean mice (Fig. 1H), we again categorized them as low- and high-runners (Fig. 2A-C). Mice suppressed their energy intake when the calorie-dilution-diet was introduced (Fig. 2D), as reported previously.(*30*) The calorie-dilution-fed mice gradually increased their daily intake from the 2^nd^ day onward until consuming more calories compared to control-fed mice on day 0 (p=0.004; Fig. 2D-G). Body weight and fat mass also decreased after introducing the calorie-dilution-diet, and they stabilized in the 2^nd^ week but without reaching the levels of control-fed mice (Fig. 2H-J). Neither absolute nor relative lean mass was influenced by calorie-dilution-diet before nor during running (Fig. S1B-D and S1G-H). During the active period, the calorie-dilution-diet groups ate more in grams but consumed a similar number of calories (Fig. 2D-G). Specifically, under calorie-dilution-diet, high-runners ate more than sedentary mice (p=0.035) and a trend was observed in low-runners (p=0.062), uncovering that energy compensation was preserved independently of the diet. However, control-fed high-runners ate more calories than calorie-dilution-fed high-runners (p=0.025), likely due to the reduction by calorie-dilution-diet of running distances (Fig. 2C,G). Indeed, the estimated energy expenditure was higher in the control-fed than calorie-dilution-fed high-runners, resulting in energy balance differences (Fig. S1I-J). Food intake was also positively associated with running distance in both diets (Fig. 2L-M). Compared to the baseline period, only control-fed high-runners significantly lost absolute and relative fat by day 35 (Fig. 2J-K and S1E-F). Together, these results reveal that energy compensation is preserved during feeding with a calorie-dilution diet.

**Fig. 2.**
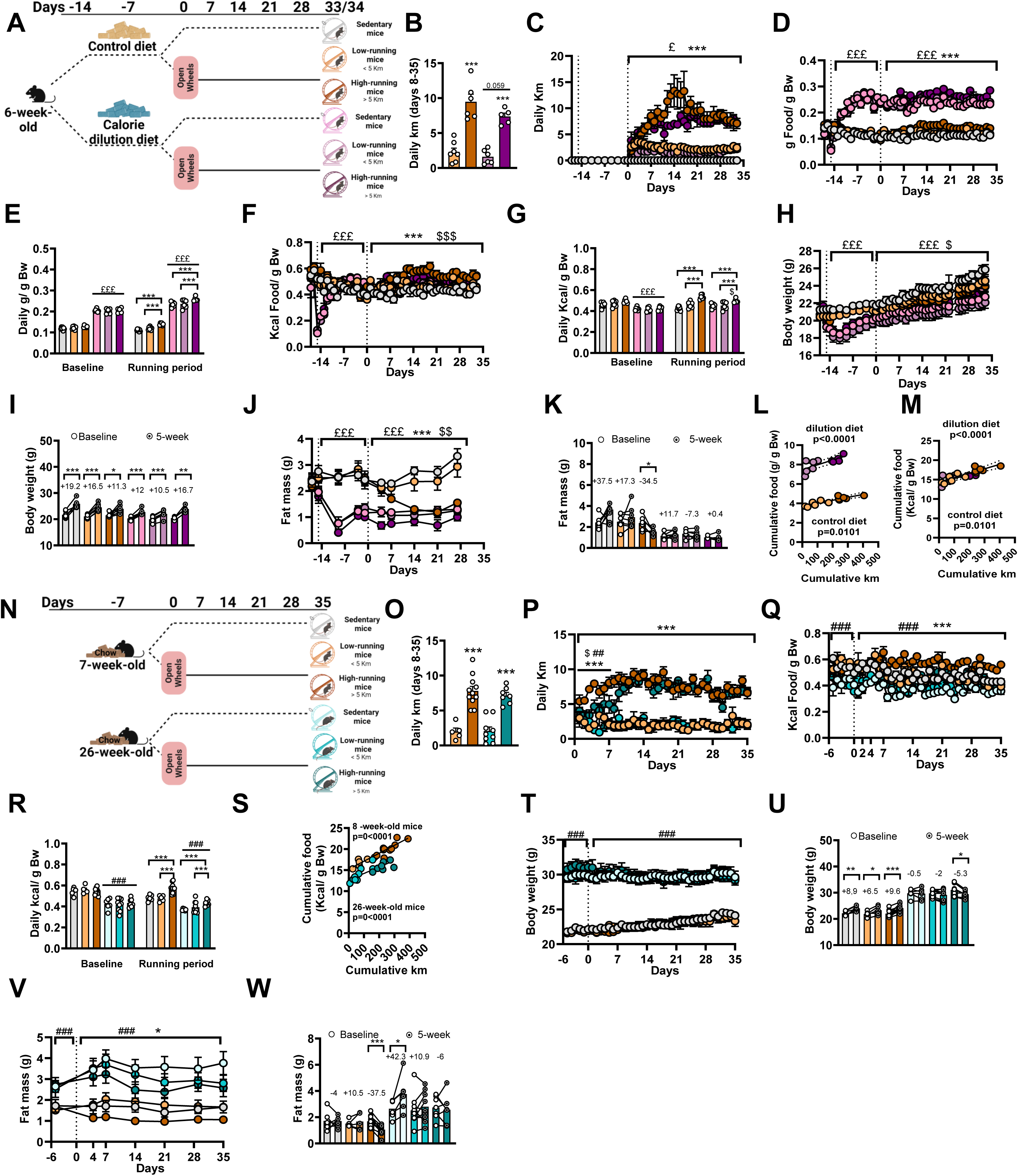
**Energy compensation is independent of diet and age. A-L**) Chronic running in young males fed a control or calorie-dilution diet**. A**, Schematic. **B**, Mean distance from day 8 to 35. **C**, Distance. **D**, Gram intake. **E**, Gram intake per period. **F**, Calorie intake. **G**, Calorie per period. **H**, Body weight. **I**, Body weight changes. **J,** Fat mass. **K**, Fat changes**. L**, Association total grams and distance. **M**, Association total calories and distance. **N-W**) Chronic running in 8- and 26-week-old mice**. N**, Schematic. **O**, Mean distance from day 8 to 35. **P**, Distance. **Q**, Calorie intake. **R**, Calorie intake per period. **S**, Association total food and distance. **T**, Body weight. **U**, Body weight changes. **V**, Fat mass. **W**, Fat changes. Two-way ANOVA with Bonferroni post hoc multiple comparison tests was used for evaluating the effects of running (*), diet (£) or age (#) and its interaction ($), but if Shapiro-Wilk’s and/or Levene’s test was significant, GzLM with sequential Bonferroni post hoc multiple comparisons was used (**B**,**E**,**G**,**O**,**R**,**U**). Two-way repeated-measures ANOVA was used to assess running (*) diet (£) or age (#) and its interaction ($), but if Mauchly’s and/or Shapiro-Wilk’s test was significant, then repeated-measures GEE was conducted (**C**,**D**,**F**,**H**,**J**,**P**,**Q**,**T**,**V**). We applied paired t test (*) in **I**,**K**,**U**,**W** and a simple linear regression in **L**,**M**,**W**. Data are mean±SEM. *P≤0.05, **P≤0.01,***P≤0.001. **A** and **N** were created using Biorender.

Next, we assessed the effect of age, which is associated with changes in body composition (*31*), on energy compensation. Moreover, to evaluate the energy compensation in a different genetic sub-strain, we exposed *ad libitum-*chow-fed young- and old-week-old C57BL/6JRj mice to wheels (Fig. 2N-P). Older mice ate less, were heavier and displayed greater fat and lean mass but similar relative fat and lower relative lean mass than younger mice (Fig. 2Q-U and S1K-Q). During the active period, older mice ran similarly to younger mice (Fig. 2O) but expended less energy and ate less than young mice, which impacted energy balance (Fig. 2O-R and S1R-S). When analyzing older mice only, high-runners ate more than sedentary mice (p=0.006) and a trend was observed in low-runners (p=0.071), highlighting that energy compensation is conserved during aging (Fig. 2Q-R). Indeed, daily food intake was positively associated to running distance in both age groups (Fig. 2S). High-running levels did not affect body weight but reduced an average of 37.5% fat and increased an average of 16.5% lean mass in younger mice, as seen above. In contrast, running led to body weight loss in older mice by blocking age-linked fat gain, resulting in an average of 6% fat loss and 4% lean loss (Fig. 2T-W and S1N-O). Upon opening the wheels, younger high-runners had an immediate drop in both absolute and relative fat mass. In contrast, older high-runners had initial increases in absolute and relative fat mass until the 2^nd^ week, at which fat mass began to decrease, coinciding with when they reached the same levels of activity as younger high-runners (Fig. 2P,V and S1O). Relative lean mass during the active period was affected by age and running, which likely resulting from differences in fat mass (Fig. S1P-Q). Together, our data suggest that energy compensation is a preserved mechanism in male mice induced by high-running levels and preceded by fat loss.

### Energy compensation is associated with circadian-specific modulation of adipose tissue and circulating leptin

To investigate how running increases food intake, we focused on the initial days of running (Fig. 3A). Runners ate less calories during the light phase on the 1^st^, 2^nd^ and 3^rd^ day of running, whereas food consumption increased in the dark phase on the 3^rd^, 4^th^ and 5^th^ day (Fig. 3B). Notably, the difference in food intake between groups disappeared during the light phase on day 4 resulting in a global increase in 24-hour food intake (Fig. S2A). Running had no effect on relative visceral and subcutaneous fat mass at ZT0 on day 3, 4 and 5 (60, 84, and 108 hours of running, respectively), which may be explained by runners eating more on the 3^rd^, 4^th^ and 5^th^ night to replenish their energy reserves. In contrast, runners reduced food intake during the light phase on day 2 and 3, which was associated with the loss of both fat mass depots at ZT12 (48 and 72 hours of running, respectively), raising the possibility that fat tissues are used as an energy source (Fig. 3C-G). Runners consumed more calories at light phase on day 4 than previous days leading to no differences between groups during this phase. Despite that, running resulted in fat mass loss of both fat depots at ZT12 on day 4 uncovering that increased food intake was not enough to compensate the energy expenditure. Running did not influence relative visceral and subcutaneous fat mass at ZT12 on day 5 (120 hours of running) highlighting that the increase in food intake had taken effect and the mouse stored the excess calories consumed as fat tissue (Fig. 3E,G).

**Fig. 3.**
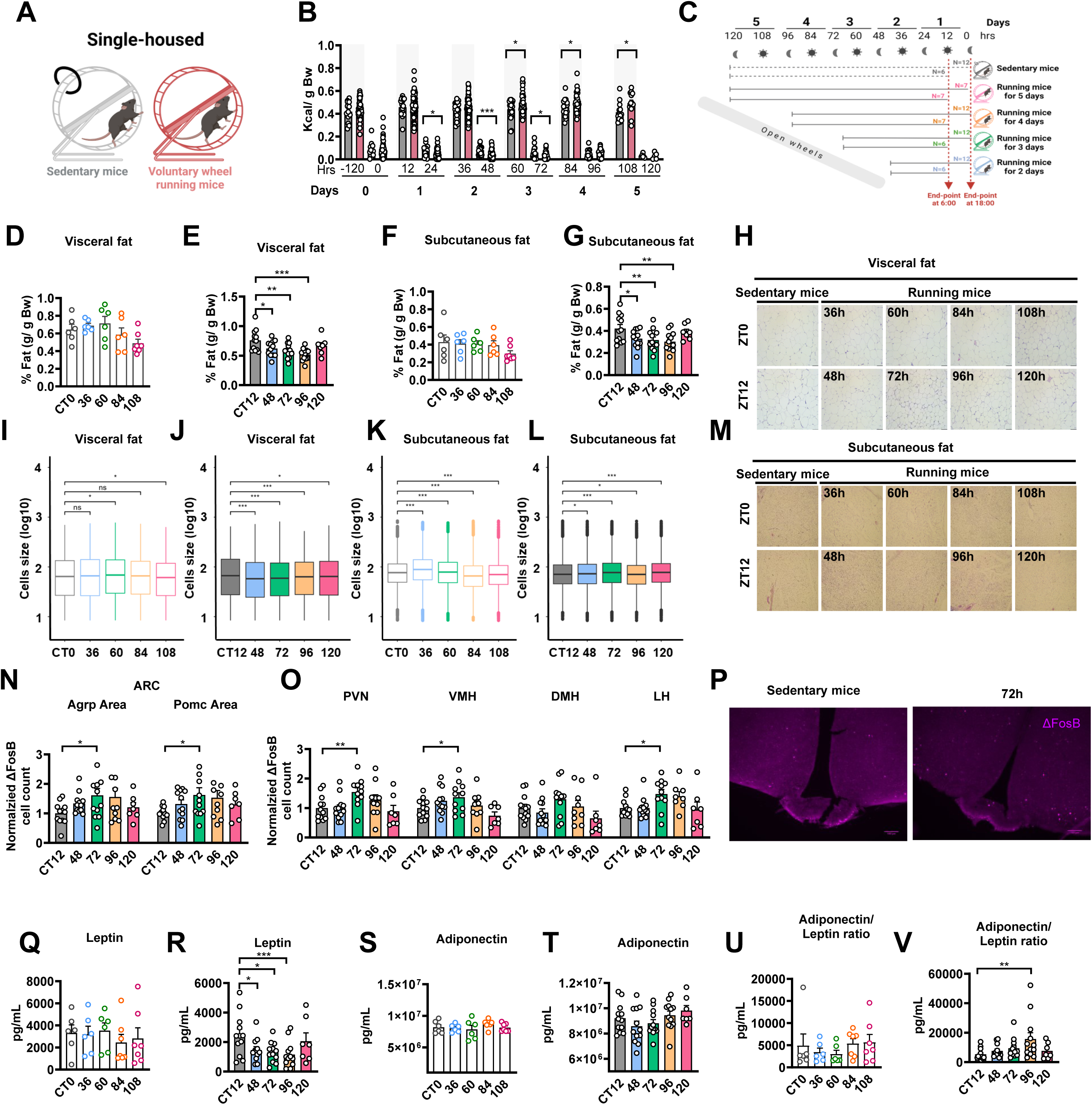
Energy compensation is preceded by simultaneously changes in fat and hypothalamus. **A**, Schematic. **B**, Calories intake. **C**, Schematic. **D**, Visceral fat at ZT0. **E**, Visceral fat at ZT12.**F**, Subcutaneous fat at ZT0. **G**, Subcutaneous fat at ZT12. **H**, Representative images of visceral fat. **I**, Visceral cell size at ZT0. **J**, Visceral cell size at ZT12. **K**, Subcutaneous cell size at ZT0. **L**, Subcutaneous cell size at ZT12. **M**, Representative images of subcutaneous fat. **N**, Chronic activation in Agrp and Pomc areas. **O**, Chronic activation in PVN, VMH, DMH and LH. **P**, Representative images of hypothalamus. Contrast was enhanced for visualization. **Q**, leptin at ZT0. **R**, leptin at ZT12. **S**, Adiponectin at ZT0. **T**, Adiponectin at ZT12. **U**, Adiponectin to leptin ratio at ZT0. **V**, Adiponectin to leptin ratio at ZT12. Data in **B** were assessed using Independent T-test for each 12-hour range and one-way ANOVA with simple contrast to reveal differences (*) regarding sedentary group was applied for **D**-**G**,**N**,**O**,**Q**-**V**. In case of non-normal distribution and/or the heterogeneous variances in the data, GzLM with simple contrast was used. Data are mean±SEM. *P≤0.05, **P≤0.01, ***P≤0.001. **A** was created using Biorender. **I**-**L** were analyzed using log10 transformed cell size groups using two-sided Mann-Whitney U tests and plotted using median.

The initiation of running had a differential impact on the fat cell size before and after energy compensation. In runners, visceral cells were larger on day 3 at ZT0, but smaller on days 2, 3, and 4 at ZT12, as well as on day 5 (Fig. 3H-J and Table S1). In contrast, subcutaneous cells were larger due to exercise on days 2 and 3, smaller on day 4 and at ZT0 on day 5 but larger at ZT12 on day5 (Fig. 3K-M and Table S1). We evaluated the molecular changes in both fat depots (Fig. S2B and Table S2). *Lrrc8Ar*, which regulates adipocyte growth (*32*), was upregulated at ZT12 on day 2 in the subcutaneous fat when t-test was used (p=0.030), confirming that the subcutaneous cells were expanded that day (Fig. S2C). The adipocyte differentiation, represented by *Peroxisome proliferator-activated receptor (PPAR)γ*, was not affected by running (Fig. S2D). However, cell proliferation in the visceral fat, represented by *Ki67* expression, was downregulated at ZT12 on days 4 and 5 (Fig. S2E). Running differentially regulated lipogenesis in a circadian rhythm-dependent manner before and after compensation. *Fatty acid synthase* (*Fasn)* in the visceral fat was downregulated at ZT0 on day 2, 3 and 4, but upregulated on day 5 at ZT12, while in the subcutaneous fat, *Fasn* was also upregulated at ZT0 on day 5 (t-test, p=0.014; Fig. S2F). Remarkably, running upregulated *β3-AR* (*Adrp3*; t-test, p=0.049), which participates in lipolysis and thermogenesis of adipocytes (*33*), in the visceral fat at ZT12 on day 3, but simultaneously decreased the Tyrosine Hydroxylase (TH; Fig. S2G-H), an essential enzyme for biosynthesis of catecholamines acting on β-ARs and regulating lipolysis (*34*). This suggests a simultaneous increased sensitivity to catecholamines and reduction in Sympathetic Nervous System (SNS) activity on that day. On the other hand, running did not influence lipolysis-related molecules, including *Lipase E, Hormone Sensitive Type* (*Lipe),* Adipose triglyceride lipase (ATGL) nor did the phosphorylation levels changes of Hormone-sensitive lipase (pHSL; Fig. S2I-K). UCP1 also remained unchanged by running initiation, suggesting that the running-induced browning of white fat is a long-term running adaptation (Fig. S2L). In addition, the phosphorylation of AMP-activated protein kinase (pAMPK), which modulates energy metabolism in fat tissue (*35*), was not affected by running initiation (Fig. S2M-N). Thus, we next quantified the circulating free fatty acids (FFA), which result from fat lipolysis and are released into the circulation to provide fuel for other tissues (Fig. S2O). Running reduced circulating FFA on day 2 at ZT12 (t-test, p=0.046), which suggests a high FFA demand as the runners still had not compensated. FFA levels were not significant on days 3 and 4 at ZT12, when overnight food intake increased. Given that no differences in running distance were observed between groups (Fig. S2P), the data indicate that energy compensation is a gradual mechanism occurring over several days in mice and associated with temporal effects on fat mass regulation.

The hypothalamus plays a central role in the regulation of whole-body energy balance. Thus, we assessed the impact of running on the hypothalamic areas by counting cFOS (acute) and ΔFosB (chronic) positive neurons at ZT12. Running increased the number of positive ΔFosB neurons in the Agrp and Pomc areas, PVN and lateral hypothalamus (LH) whereas it increased both acute and chronic activation of the ventromedial hypothalamus (VMH) neurons on day 3 (Fig. 3N-P and S3A-B). It has been described that chronic running activates the dorsomedial portion of the VMH (dmVMH) neurons, which in turn, also express the leptin receptor.(*36*) While leptin receptors are expressed in different hypothalamic nuclei (*14*, *15*), their primary location within the VMH is the dmVMH (*37*). Here, running acutely activated the dmVMH neurons (t-test, p=0.048) and decreased Agrp intensity in the VMH on day 3 (t-test, p=0.038; Fig. S3C-E). Therefore, we next analyzed circulating leptin. While no difference in circulating leptin was observed between groups at ZT0 (Fig. 3Q), runners declined leptin at ZT12 on days 2, 3, and 4 whereas increased it on day 5 (Fig. 3R), possibly due to increased food intake and fat mass (*23*). To validate the increase in leptin after compensation, we analyzed circulating leptin in 5-week (chronic) running mice and, as expected, no differences were observed at ZT0 or ZT12 between sedentary and running mice (Fig. S3F). Remarkably, neither significant alterations in *leptin* expression nor protein levels were observed in any fat depot between groups after running initiation (Fig. S3G-J). Typically, leptin decline is associated to a reduction in fat stores (*38*). Thus, we explored whether circadian-specific modulation of circulating leptin correlate with fat mass. In sedentary mice, leptin correlated positively with both fat depots at ZT12, but not at ZT0. In runners, circulating leptin at ZT12 was positively associated with relative visceral fat on days 4 and 5, and with subcutaneous fat on day 5, with a trend on day 4. At ZT0, leptin correlated positively with visceral fat on days 2, 3, and 4, with a trend on day 5, and with subcutaneous fat on days 2 and 5, and a trend on day 4 (Fig. S3K). Moreover, mice with lower circulating leptin also exhibited lower FFA levels at ZT12 on days 3 and 4 (Fig. S3L). In summary, these results underpin that running regulates leptin in a circadian-specific manner before energy compensation.

Adiponectin increases food intake, opposing leptin’s role in the hypothalamus (*24*). Here, *adiponectin* in subcutaneous fat was upregulated on day 4, particularly at ZT0 (Fig. S3G-H), although the tissular protein levels remained unchanged (Fig. S3I-J). Circulating adiponectin slightly declined on days 2 and 3 reaching sedentary levels on day 4 and 5 at ZT12, resulting in a significant adiponectin to leptin ratio difference between sedentary mice and runners only on day 4 at ZT12 (Fig. 3S-V). This change occurred simultaneously with the increase in 24-hour food intake by running (Fig. 3B), which uncover that modulation of adiponectin may contribute energy compensation.

### The β3-AR mediates running-induced fat loss and the lower leptin triggers energy compensation

Lipolysis is mediated in part by the β-adrenergic system (*26*). Since β3-AR was upregulated in the visceral fat before compensation (Fig. S2G), the β3-AR might mediate fat mobilization, thus leading to energy compensation. Therefore, we injected mice with β3-AR antagonist SR52930a or vehicle control and provided wheels access. In addition, half of the runners injected with the vehicle were given SR52930a from day 3 of running (3d-SR52930a-runners) to evaluate whether SR52930a affects distance and indirectly impacted on fat and food intake (Fig. 4A). SR52930a did not alter cumulative distance or food intake before running or from day 3 (Fig. 4B-D). Remarkably, there was an association between total distance and total food in vehicle-runners and 3d-SR52930a-runners, but not in SR52930a-runners, suggesting that the blockage of β3-AR before running regulates energy compensation (Fig. 4E). No overall body weight changes were observed between groups, but SR52930a prevented absolute fat loss before running (Fig. 4F). Only vehicle- and 3d-SR52930a-runners displayed a significant lean gain over the experiment (Fig. 4G). Our findings emphasize the importance of intact β3-AR activity for mediating running-induced fat loss and subsequent energy compensation.

**Fig. 4.**
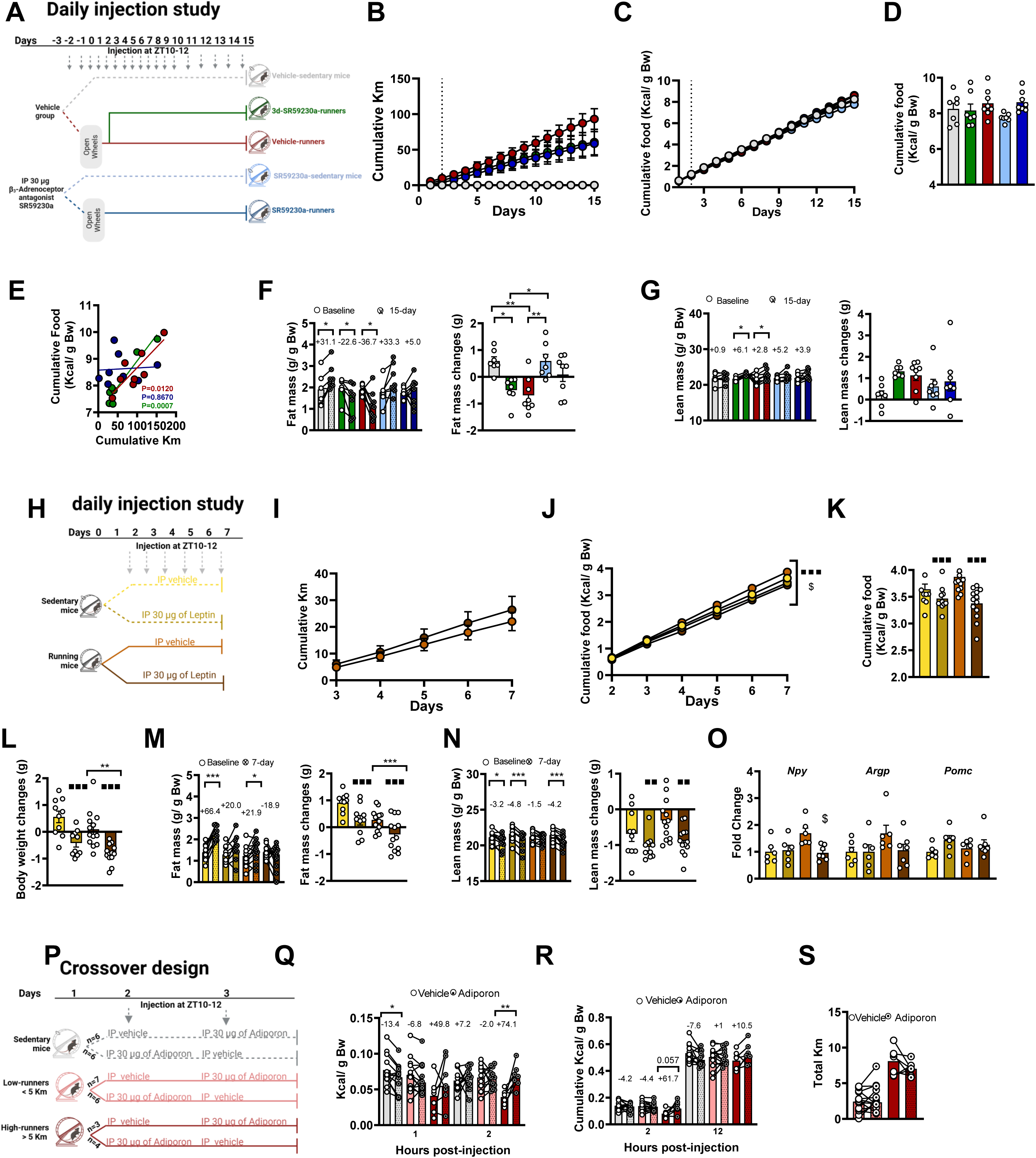
β3-AR drives fat loss and circulating decline of leptin induces energy compensation. **A**-**G**) Daily SR52930a injection study. **A**, Schematic. **B**, Total distance. **C**, Total intake. **D**, Total intake after 15 days. **E**, Association total intake and distance. **F**, Fat changes. **G**, Lean changes. **H-N**) Daily leptin injection study**. H**, Schematic. **I**, Total distance. **J**, Total intake. **K**, Total intake after 7 days. **L**, Body weight changes. **M**, Fat changes. **N**, Lean changes. **O**, Hypothalamic gene expression **P-S**) Crossover design with Adiporon. **P**, Schematic. **Q**, Hourly food intake. **R**, Cumulative food. **S**, Total distance. One-way repeated-measures ANOVA was used to assess the running (*) in **B**,**C**,**I** whereas two-way repeated-measures ANOVA was used to analyze the running (*), drug (′) and its interaction ($) in **J**. But, in both cases, whether sample’s sphericity and/or normality test were significant, repeated-measures GEE was used. One-way ANOVA with Bonferroni post hoc multiple comparison tests was used to evaluate the running (*) in **D**,**F**,**G** whereas two-way ANOVA with Bonferroni post hoc multiple comparison tests was used to evaluate the running (*), drug (′) and its interaction ($) in **K**-**O.** However, if data did not follow a normal distribution and/or the variances were not homogeneous, GzLM with sequential Bonferroni post hoc multiple comparisons was used. Paired T-test (*) was applied in **F**,**G**,**M,N**,**Q**-**R** and simple linear regression was used in **E**. Data are mean±SEM. *P≤0.05,**P≤0.01, ***P≤0.001. **A**, **H** and **P** was created using Biorender.

Since β3-AR blockade prevented running-induced fat loss (Fig. 4F) and β3-AR stimulation is known to suppress *leptin* in white fat (*28*), we hypothesized that leptin lowering might be a major driver of energy compensation, as its levels declined before energy compensation (Fig. 3R). Therefore, mice were injected daily with leptin or vehicle from day 1 to ensure equal phenotypic distribution (Fig. 4H). We used a low dose of leptin to avoid changes in food intake in sedentary mice. Leptin injection did not affect running distance but reduced food intake in runners (t-test, p=0.0003) and not in sedentary mice (t-test, p=0.178; Fig. 4I-K). On day 7, leptin reduced body weight and composition whereas running decreased body weight and fat mass (Fig. 4L-N). However, relative fat changes persisted due to leptin whereas leptin’s effect on relative lean mass changes were smaller, indicating at crucial role of leptin predominantly on fat regulation (Fig. S4A-C). Next, we explored the hypothalamic changes on day 3 in another cohort. Running upregulated *Npy*, but this effect was blunted by leptin, and a similar trend was observed for *Agrp*. Leptin slightly upregulated *Pomc* (t-test; p=0.054) without overall changes regarding running (Fig. 4O). Leptin regulates *Steroidogenic factor-1* (*Sf1)*-VMH neurons to control body weight and suppress food intake via hypothalamic Basonuclin-2 (Bcn2)-neurons (*39*, *40*). Here, neither *Bcn2*, *Nr5a1*, which encodes for SF-1, nor *Leptin receptor* were altered by running or leptin (Fig. S4D). Together, we reveal that circulating leptin decline drives energy compensation after running initiation by upregulating *Npy* expression.

Adiponectin promotes food intake under fasting conditions and reverses central leptin-induced suppression of food intake (*24*). Here we observed a simultaneous increase in 24-hour food intake and adiponectin to leptin ratio on day 4, and a downregulation of hypothalamic *AdipoR1* by running on day 3 (Fig. 3V, S2A and S4L). To assess whether adiponectin impacts on food intake in high-runners with negative energy balance, mice were injected with an adiponectin agonist receptor, Adiporon, in a paired design experiment (Fig. 4P). Before compensation, high-runners ate less than sedentary mice and low-runners in the first two hours, which is likely due to increased physical activity. Adiporon decreased food intake by an average of 13.4% in sedentary mice during the 1^st^ hour, while it increased food intake in high-runners by an average of 49.8% and 74.1% during the 1^st^ and 2^nd^ hours, respectively, reaching levels similar to sedentary mice by the 2^nd^ hour (Fig. 4Q). This resulted in a cumulative increase in food intake of an average of 61.7% and 10.5% at 2- and 12-hour post-injection, respectively, without effecting total running distance (Fig. 4R-S). Adiporon did not affect food intake in high-runners after energy compensation (Fig. S4E-G). Overall, we conclude that activation of β3-AR drives running-derived fat loss and leptin decline triggers running-derived energy compensation. Moreover, high-runners are potentially sensitive to adiponectin only under negative energy balance.

### Running induces a new homeostatic steady state in young mice

To explore the effects of running in the new baseline state, we blocked the wheels from younger and older mice of Fig. 2N. Food intake of the now inactive-high-runners gradually decreased, fluctuating through the days, although they still ate more than inactive-low-runners and sedentary mice (Fig. 5A-C). There were no differences in body weight and lean mass between groups, but both absolute and relative fat of younger inactive-high-runners slightly increased whereas those from older inactive-high-runners reached same levels as age-matched sedentary mice (Fig. 5D-E and S5A-E). Indeed, younger inactive-high-runners showed a 29.8% fat loss whereas older inactive-high-runners displayed a 30.9% fat gain by day 45 (Fig. 5F). While the physiological parameters in younger sedentary mice did not display any association, older ones showed a trend towards a negative association between fat on day 35 and food intake during inactivity. This pattern was further accentuated in both inactive groups, which also showed a positive association of running distance with food intake. However, only older inactive-mice exhibited a positive association between food intake and fat gain, and between running distance and fat gain (Fig. S5F). This uncovers that fat is regulated in an age-dependent manner during inactivity. As expected, lean mass on day 35 was not associated with food intake in any of the groups (Fig. S5G). Energy expenditure abruptly decreased on the first day in younger inactive-high-runners and slowly in older inactive-high-runners yielding no difference in energy balance between groups (Fig. S5H-J). These data suggest that energy compensation persists during inactivity in both young and old mice, but it is associated with lower fat mass in young mice and a rebound in fat mass in old mice.

**Fig. 5.**
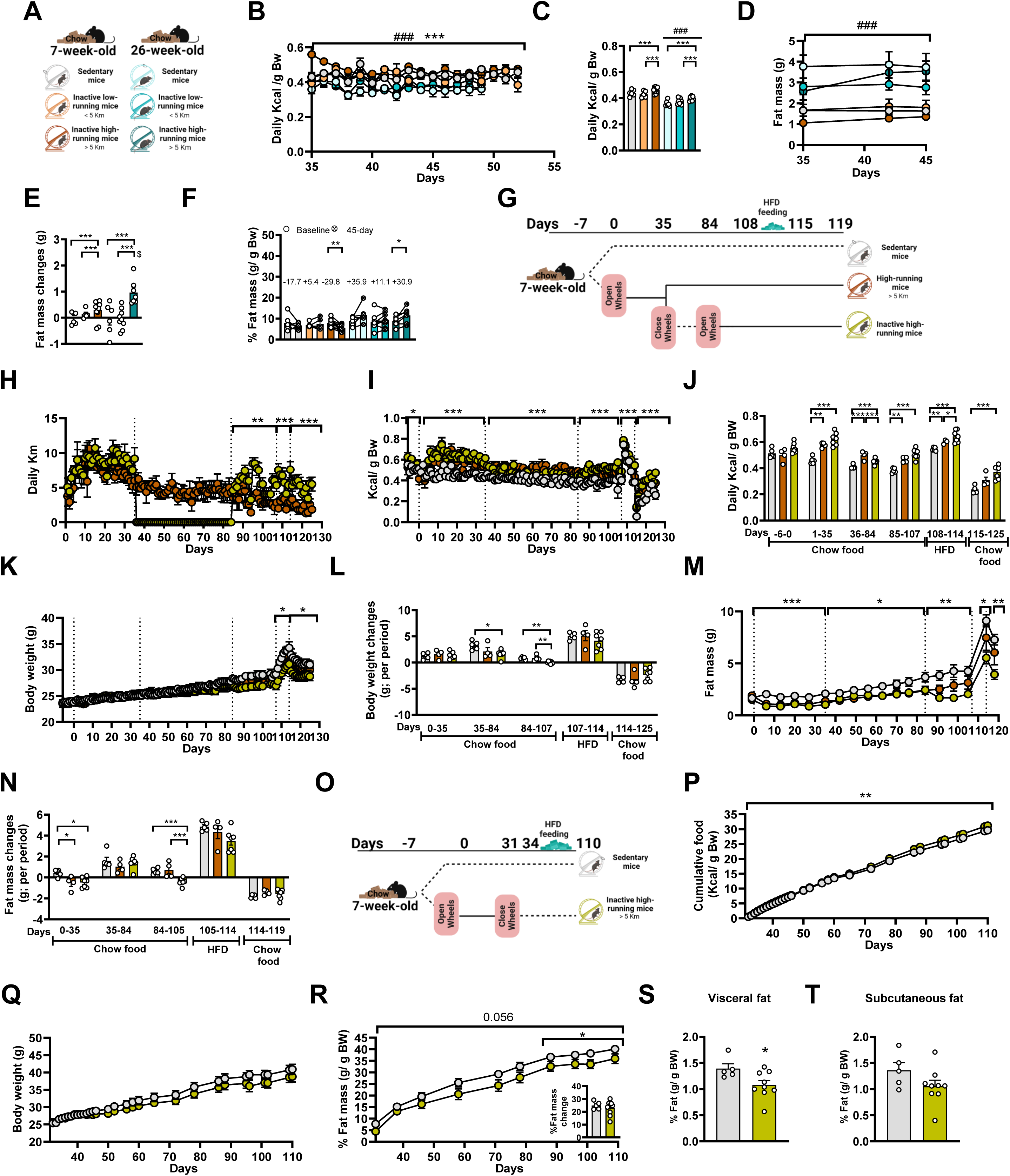
**Running induces a new steady state. A-F**) Effect of age during inactivity. **A**, Schematic. **B**, Food intake. **C**, Intake during period. **D**, Fat mass. **E**, Fat changes. **F**, Relative fat mass. **G**-**N**) Effect of long-term inactivity. **G**, Schematic. **H**, Distance. **I**, Intake. **J**, Intake per period. **K**, Body weight. **L**, Body weight changes. **M**, Fat mass. **N**, Fat changes. **O**-**T**) Effect of HFD feeding during inactivity. **O**, Schematic. **P**, Total intake. **Q**, Body weight. **R**, Relative fat mass. **S**. Relative visceral fat. **T**, Relative subcutaneous fat. One-way repeated-measures ANOVA was conducted to evaluate the main effect of running (*) in **H**,**I**,**K**,**M**,**P**,**Q**,**R** and Two-way repeated-measures ANOVA was used to assess the effect of running (*), age (#) and its interaction ($) in **B**,**D**. In case that sample’s sphericity and/or sample’s normality test was significant, repeated-measures GEE was used. One-way ANOVA was conducted to evaluate the main effect of running (*) in **J**,**L**,**N** and two-way ANOVA was conducted to evaluate the main effect of running (*), age (#) and its interaction ($) in **C**,**E**. In case of non-normal distribution and/or heterogenous variances, GzLM with sequential Bonferroni post hoc multiple comparisons was used. Independent t-student was applied to assess differences between group in **R**,**S**,**T** whereas paired t-test was used for evaluating difference for each group in **F**. Data are mean±SEM. *P≤0.05 **P≤0.01, ***P≤0.001. **A**, **G** and **O** was created using Biorender.

We then examined the metabolic phenotype in another young cohort during prolonged inactivity, followed by a 2^nd^ active period with the challenge of a HFD (Fig. 5G-H). Food intake fluctuated over the long-term inactive groups (day 36 to 84), where chronic-high-runners ate the most, followed by inactive-high-runners and lastly sedentary mice (Fig. 5I-J). On day 84, inactive-high-runners were lighter than sedentary mice, likely caused by lean loss (Fig. 5K-L and S5K-M). Absolute and relative fat differences on day 35 between groups persisted throughout the inactivity due to similar fat gain (Fig. 5M-N and S5N). After reopening the wheels (day 85 to 107), inactive-high-runners lost fat mass, which prevented body weight gain despite the food intake increase (Fig. 5K-N). The inactive-high-runners ran more than the chronic-high-runners during the 2^nd^ active period, suggesting that differences in body weight and fat mass may stem from changes in activity status and/or running distance (Fig. 5H). Next, we assessed food preferences by providing them simultaneously a HFD and chow food on days 108-109. The three groups ate more calories from the HFD, but running mitigated HFD-induced hyperphagia and controlled overconsumption throughout the week (Fig. S5O-R). Differences in food intake between running groups likely resulted from differences in their traveled distance (Fig. 5H-J). Subsequently, the HFD withdrawal suppressed chow intake (Fig. S5S), which has been previously described.(*41*) However, we show here for the first time that chow devaluation was higher in sedentary mice than inactive-high-runners both on the first and last days of chow exposure, and interestingly, chow devaluation negatively correlates with distance on the last day (Fig. S5S-T). Although no differences in body weight changes were observed during HFD exposure and withdrawal (Fig. 5L), slight differences in absolute fat mass were observed between inactive-high runners and sedentary mice due to HFD exposure (Fig. 5N), accentuating the initial differences between groups (Fig. 5M).

Finally, we assessed the metabolic phenotype of inactive-high-runners by exposing a new cohort to long-term HFD-feeding during inactivity. Mice were given both HFD and chow to evaluate food preferences on days 3 and 4 of inactivity, then were exclusively HFD-fed (Fig. 5O). Both inactive-high-runners and sedentary mice preferred HFD and no differences between groups were observed during the 3^rd^ day of inactivity, suggesting that inactive-high-runners may control better HFD-hyperphagia of 1^st^ day of exposition (Fig. S5U). However, inactive-high-runners ate more HFD on the 2^nd^ day and over the experiment (Fig. 5P and S5U), highlighting that the feeding pattern of running persists during inactivity. Although no changes were observed in body weight, the relative fat mass of inactive high-runners tended to be lower than sedentary mice over time, being more pronounced during the plateau period (Fig. 5Q-R). Inactive-high-runners were less viscerally obese than sedentary mice and same tend was observed for subcutaneous fat (Fig. 5S-T). Together, the results evidence that running in youth protects from fat mass rebound during inactivity in both lean and obese mice.

### Chronic running modulates hypothalamic hunger signaling with clinically relevant effects

Given the surprising impact of chronic running on food intake in the new hemostatic set point, we analyzed hypothalamic adaptations to chronic running and inactivity using bulk RNA-sequencing (Fig. 6A). The PCA showed a minimal clustering between groups (Fig. 6B). Most of overrepresented biological GO processes between sedentary mice and high-runners were related to biogenesis and assembly of ribonucleoprotein structures whereas those between sedentary and inactive mice were linked to synapsis (Fig. 6C). However, some DE genes of inactive mice vs high-runners comparison were associated to feeding behavior. Indeed, *Npy* was upregulated by running, whereas *leptin receptor* was upregulated by inactivity (Fig. 6C, S6A and Table S3-5). Interestingly, *Socs3*, a leptin-regulated inhibitor, was downregulated by running whereas *Mapk3* was upregulated by running and inactivity (Table S3). *Mapk3* encodes ERK1, which mediates the hypothalamic leptin effects and influences glucose-regulated *Pomc* expression (*42–44*). These results uncover that chronic running may induce molecular hypothalamic adaptations in the leptin signaling, which may persist during new steady state.

**Fig. 6.**
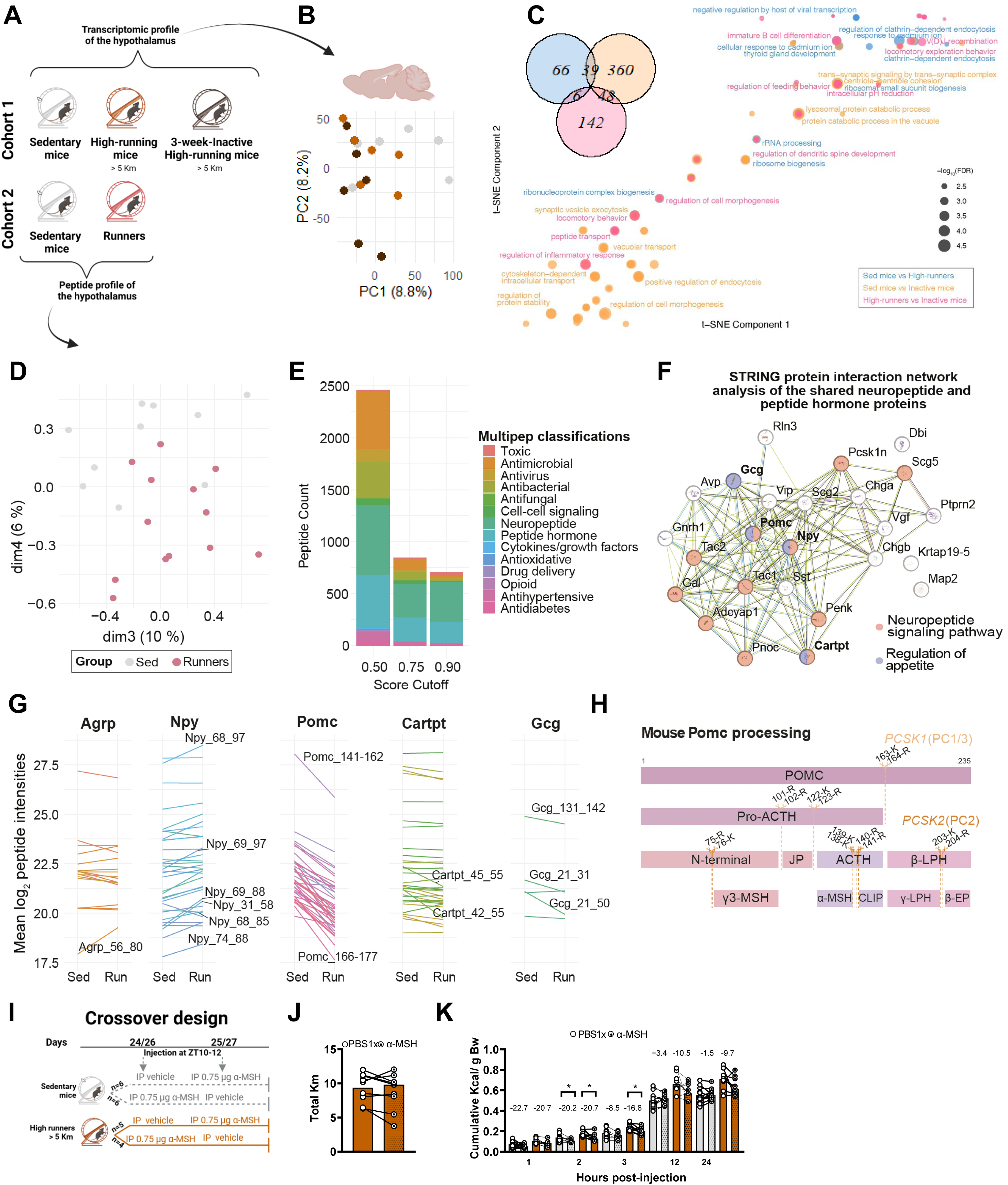
Hypothalamic adaptations. **A**, Schematic. **B**, Transcriptomic PCA. **C**, GO biological process enrichments. **D**, Peptidomics PCA. **E**, Bioactivity classification. **F**, String protein interaction network analysis of the shared neuropeptide and peptide hormone proteins. **G**, Bar plots showing the mean log2 intensities of peptides. **H**, *Pomc* processing of rodent hypothalamus. **I-K**) Crossover design with α-MSH short cyclic analogue. **I**, Schematic. **J**, Distance. **K**, Cumulative intake. Data are mean ± SEM. *P ≤ 0.05 **P ≤ 0.01, ***P ≤ 0.001. **A** and **I** was created using Biorender. Paired T-test was applied in **J** and **K** for assessing the effect of each group.

Finally, we performed a peptidomics analysis on the hypothalamus of another cohort of sedentary mice and runners to elucidate the effects of running on hunger signaling mediated by peptides. PCA enabled separation between groups (Fig. 6D), highlighting that variance in peptide levels is driven by running status. Identified peptide bioactivity was mainly attributed to the Neuropeptide and Peptide Hormone classifications (Fig. 6E). These two categories shared 78 (out of 585) peptides, corresponding to 25 proteins with neuropeptide signaling and appetite functions (Fig. 6F). Running tended to increase the abundance of Agrp_56-80, Npy_68-97, Npy_69-97, Npy_69-88, Npy_31-58 and Npy_74-88 whereas it tended to reduce the abundance of all Pomc peptides, Cartpt_42-55, Cartpt_45-55, Ggc_131-142, Ggc_21-31 and Ggc_21-50 (Fig. 6G and Table S6). The consistent tendency to reduce the abundance of all Pomc peptides underlines that running alters *Pomc* processing (Fig. 6H). In fact, we detected various cleaved fragments of Pomc, including 4 N-terminal (sequences 51-75), 7 joining peptide (JP; sequences 103-120), 7 corticotropin-like intermediate peptide (CLIP; sequences 141-162), 26 γ-lipotropin (γ-LPH; sequences 165-195) and 2 β-endorphin (β-EP; sequences 209-222). Of note, γ-LPH and β-EP inhibits and stimulates food intake, respectively (*45*, *46*). Food intake was significantly and positively correlated with Gal_33-45 and Npy_68-85 in sedentary mice and Arf1_160-181 and Penk-218_227 in runners (Fig. S6B). While Galanin and NPY stimulate feeding,(*47*, *48*) the role of Arf1 and Penk in regulating food intake remain unknown. The collapsed data identified a total of 1504 unique proteins for GO enrichment, retrieving only overrepresented molecular function terms (Fig. S7C and Table S6). Cathepsin L and Pcsk2 were listed in the catalytic activity term, and both are related to *Pomc* processing (*49*, *50*). Despite that proteolytic processing of *Pomc* produces α-MSH (*49*), we did not detect in the peptidome. However, we speculated that running may enhance α-MSH sensitivity whether *Pomc* processing is altering by running. Thus, sedentary and running mice were injected with α-MSH analogue in a paired design experiment (Fig. 6I). The effect of α-MSH on food intake was prolonged in high-runners than sedentary mice without changes in running distance (Fig. 6J-K). Together, the data reveal that running induces hypothalamic adaptations in hunger signaling with promising clinical consequences.

## DISCUSSION

Here, we report that running improves the coupling between energy intake and expenditure, leading to favorable changes in body composition without significant alterations in body weight, which is attributable to energy compensation. Running-induced energy compensation was a response that depended on initial distance, was preceded by SNS-mediated fat loss and circadian-dependent decline in circulating leptin, and persisted even after cessation of active period. Leptin replacement prevented this mechanism and blocked *Npy* upregulation by running, uncovering that decreased leptin is required for the compensatory increase in food intake, consistent with leptin’s physiological role.

Our molecular results suggest that energy compensation is a time-dependent gradual mechanism coordinated by utilization of both visceral and subcutaneous fat depots. This aligns with the lipostat theory, which postulates that fat-derived factors act on the brain to regulate energy balance (*51*). The blockage of β3-AR before running prevented fat loss and altered the food-distance association. Considering the hierarchical role of SNS in fat utilization during calorie restriction (*52*), further research is needed to address the specific role of β3-AR in each fat depot during running. Here, running aggravated the circulating leptin decline at ZT12, further accentuating its lowest point during the daily oscillation (*53*), and the decline of circulating leptin was identified as a trigger of energy compensation. On day 3 of running, we observed chronic and acute activation of VMH, mainly acute in the dmVMH, and a chronic transiently activation of ARC, PVN and LH. While other running-derived factors may influence these areas (*54*, *55*), it is likely that part of this activation is mediated by leptin, especially in the ARC and dmVMH (*14*, *15*, *36*, *40*). Indeed, further analysis revealed that leptin replacement blocks *Npy* upregulation by running on that day.

Running-induced energy compensation was preserved across different diets, ages and sub-strains of lean mice. The calorie-dilution-diet also reduced running distances, which may explain the decreased energy expenditure. In contrast, age reduced the energy expenditure of low- and high-runners without affecting the average running distance, emphasizing that other compensatory mechanisms may regulate energy demand during running in older mice. Indeed, increased off-wheel inactivity was observed in 3-6-month-old mice (*56*), which may also explain the plateau phase in energy expenditure in young chow-fed high-runners after 1^st^ week, despite slight increases in running thereafter. Beyond increased energy expenditure and intake, it is known that running attenuates HFD-hyperphagia and reverts the HFD-induced alterations in food preference (*11*, *57*, *58*). Here, we were surprised that running also protected against HFD-related chow devaluation, uncovering an improved adaption to diet change. Chow devaluation, even in states of caloric deprivation, is encoded by *Agrp*-expressing neurons and mesolimbic dopamine pathway (*41*). Whether running modifies these neural basis warrants future investigation.

Chronic running induced a new metabolic steady state in young mice, where increased energy expenditure lasted only few days, as previously reported (*36*), but increased food intake persisted after running without a rebound in fat mass in lean and obese young males. Enhanced leptin sensitivity is a beneficial effect of chronic running (*59*, *60*). We speculate, based on our results, that this may persist in the new steady state. Whereas chronic exercise increases leptin sensitivity in lean conditions (*59*), maybe through the orexin neurons activation,(*60*) increased leptin receptor binding in VMH and DMH and pSTAT3 in the ARC have been described as underlying mechanisms of exercise protection against obesity (*36*, *61*). Chronic activation of dmVMH neurons, which occurred independently of fat mass, is another proposed leptin sensitizing mechanism during chronic running (*36*). Here, the chronic activity of Agrp and Pomc areas was similar to sedentary conditions on both the compensatory day (day4) and the day after, whereas the activation of dmVMH neurons decreased the next day of energy compensation (day 5; t-test, p=0.036), likely due to an increase in circulating leptin. *Npy* was upregulated by running before food compensation, and, after energy compensation, chronic running trended to increase the abundance of Npy and Agrp peptides but decrease Pomc peptides, aligning with previous results in which running downregulated *Pomc* expression assessed by *in situ* hybridization (*62*). Running remodels synaptic and cellular properties of Npy- and Pomc-neurons, with *leptin receptor-*expressing Pomc-neurons being more sensitive (*63*). Thus, it is possible that the leptin sensitivity of these neurons increases over time as part of the energy compensation system during running, potentially contributing to the new metabolic homeostasis state after the cessation of the active period. Indeed, both chronic-high-running and inactivity affected hypothalamic genes involved in leptin signaling. Running also increased sensitivity to α- MSH agonist, a proteolytic fragment of *Pomc*, supporting the idea that running induces adaptations in hypothalamic neuropeptide signaling, which modulates feeding behavior and energy balance.

In conclusion, our study unifies and extend numerous previous studies demonstrating that running-induced energy compensation is regulated by body energy stores and circulating leptin, supporting the lipostat theory of energy homeostasis. Notable, β3-AR activity modulates running-induced fat loss, while leptin replacement inhibits energy compensation and blocks running-derived *Npy* upregulation, suggesting reduced leptin as a driver of this process. We also observed that chronic running not only sustains increased food intake and improves regulation of HFD-related hyperphagia, but also protects against HFD-related chow devaluation and was associated with hypothalamic hunger signaling adaptations. Interestingly, the increased food intake induced by running persisted during inactivity and is associated with lower fat mass in lean and obese young mice. These findings uncover a new homeostatic steady state driven by running in young mice.

### Limitations of the study

Voluntary wheel running was used to avoid the activation of stress-related brain areas by treadmill running (*64*). Thus, running was an uncontrolled variable. All mice were chronically single-housed, which we showed that affect body weight and composition, thereby each cohort had different baseline periods until physiological parameters were normalized. Although transcriptome and peptidome data showed minor changes after applying the corrected p-value, we explored the biological mechanisms and overall data still suggest that running may induce hypothalamic adaptations. We did not examine the effect of sex, which is known to impact appetite regulation in humans (*65*), or pathological conditions.

## METHODS

### Ethical statement

Animal experiments complied with the European directives regarding the care and use of experimental animals and approved by the Danish Animal Experiments Inspectorate (2020-15-0201-00599). C57BL/6NRj mice were purchased from Janvier Labs and maintained at the animal facilities of the University of Copenhagen, Denmark. In case of study 3 (see below) C57BL/6JRj (Janvier Labs) were used. All mice were 7/8-week-old upon arrival, unless otherwise indicated.

### Mice and Experimental diets

All animals used in this study were housed at 22±2C with a 12-h light/dark circle (light on at 6AM (ZT0) and light off at 6PM (ZT12) with a preceding 30 minutes of half-light in each period). Mice were provided with nesting, bedding material, small plastic house and cardboard tube. Male mice were single-housed and had *ad libitum* access to water. Mice were regularly feed *ad libitum* chow diet (SAFE D30, Safe Diets, 3.389 Kcal/gram; 26 % energy from protein, 14.1 % energy from fat and 60% energy from carbohydrates; Europe, www.safe-lab.com), or when indicated to *ad libitum* high-fat diet (HFD) (D12492, 5.24 kcal; 20% energy from protein, 60% energy from fat and 20% energy from carbohydrates), control diet of calorie dilution diet (D12450B, 3.8 Kcal/gram; 20% energy from protein, 10% energy from fat and 70% energy from carbohydrates) and calorie dilution diet (D16061505, 20% energy from protein, 10% energy from fat and 70% energy from carbohydrates. In the latter, 50 % cellulose has been added, and the diet holds 1.9 kcal/g). Both HFD, calorie dilution diet and or control diet of calorie dilution diet were purchased from Research Diets Inc, USA (www.researchdiets.com).

### Experiments and diets

Body weight and energy intake were measured daily before the light off. The wheels were always opened in all studies before the light off counting the first night of running as day 1 of experiment. Body fat mass and lean mass were quantified weekly by an EchoMRI Body Composition Analyzer. In all cases, mice were divided into experimental group matching for body weight, body composition and food intake. Days of baseline differed between experiments since we waited until food intake and body weight were stabilized. Specific datapoints were excluded both for sedentary and running mice owing to measurement errors, then mean from day before and after were used. In case of daily measurements could not be taken, last and first day of missing data was averaged. Several datapoints were also missed owing to technical problems with the counters.

Statistical analyses were performed in IBM SPSS Statistics v29.0.1.0. Statistical analysis information for each graph can be found in the Fig. legends. No statistical methods were applied to predetermine the sample size for experiments. All data are shown as mean ± SEM, and findings with *P ≤ 0.05, **P ≤ 0.01, ***P ≤ 0.001 were considered statistically significant.

### Study 1: Effect of voluntary wheel running on body weight, body composition and food intake

Two young cohorts of mice were conducted independently and plotted together: study 1.1 (12 sedentary and 12 running mice) and study 1.2 (10 sedentary and 14 running mice) after 20 and 9 days of baseline respectively. Measurements of study 1.1 were annotated daily whereas measurements of study 1.2 were annotated daily until day 7, afterwards data was calculated using the average between different days. Mice were scanned during baseline period several times to habituate them to be scanned. All mice were scanned on day 2,4,7,21 and 28. However, mice from study 1.1 were scanned on day 14 whereas those from study 1.2. were scanned on day 13 but data was plotted together representing changes at 2^nd^ week of running. Study 1.2. was finished on day 35 to determine the impact on exercise on 5-week running at ZT0 whereas mice from study 1.3 were euthanized and hypothalamus were used for neuropeptide analysis.

### Study 2: Effect of calorie dilution diet on physiological response to exercise

Thirty-eight 4-week-old mice were ordered and single-housed and fed with control diet immediately when arrived. After four days and based on the equal distribution of mice regarding body composition and food intake, 18 mice were fed with calorie dilution diet. After 15 days, 7 control-fed and 7 dilution diet-fed mice were provided with blocked wheels and 13 control-fed and 11 dilution diet-fed mice were provided with functional for 33 and 34 days. Tissues have not been analyzed.

### Study 3: Effect of age on physiological response to exercise and inactivity

Twenty-two 7-week-old mice and twenty-two 26-week-old mice were ordered and single-housed immediately when arrived. After 12-days and based on the equal distribution of mice regarding body composition and food intake, 6 7-week-old and 6 26-week-old mice were provided with blocked wheels and 16 7-week-old, and 16 26-week-old mice were provided with functional for 35 days. On day 35, functional wheels were closed, and 7-week-old mice were examined for 17 days whereas 26-week-old mice were assessed for 14 days. Tissues have not been analyzed.

### Study 4: Effect of running on food intake and analysis of the underlying mechanism

The study was performed twice, and groups were pooled: study 5.1 (5 sedentary mice, 5 48 hour-running mice, 5 72 hour-running mice and 5 96 hour-running mice) and study 5.2 (13 sedentary mice, 6 36 hour-running mice, 7 48 hour-running mice, 6 60 hour-running mice, 7 72 hour-running mice, 7 84 hour-running mice, 7 96 hour-running mice, 7 108 hour-running mice and 7 120-hour running mice). In both experiments, mice were divided into sedentary or different running groups based on matched daily baseline body weight and food intake and wheels were opened in different days to sacrifice the mice the same day avoiding day-effect impact on the outcomes. In both cases, the wheels of first group of running were opened after 16 days of baseline. All mice from study 5.1 and 7 sedentary mice and 48-,72-,96- and 120-running mice from study 5.2 were sacrificed after the lights was on (ZT12). In contrast, 6 sedentary mice and 36-, 60-, 84-, and 108-hour-running mice were sacrificed after the lights was on (ZT0). Mice from study 5.2 were sacrificed the same day.

### Study 5: Physiological response of high-runners to long-term inactivity period

Male mice were single- housed immediately after arrival and fed with a chow *ad libitum* diet. Twenty days after arriving, mice were divided into sedentary with blocked wheels (12 mice) and running mice with functional wheels for 35 days (12 mice). On day 35, running mice were divided in two group: 4 mice that kept functional wheels (chronic active mice) and 8 mice that wheels were closed for 49 days (inactive mice). This was conducted based on equal distribution of food intake and fat mass on day 35. After this inactivity period, we opened the wheels from 8 inactive mice and 7 sedentary mice (new active mice). Since new active mice were run a mean of 2.03±0.81 Km they were excluded of data analysis since no differences were observed regarding bodily measurements and food intake compared to sedentary mice, as expected. Mice were fed with chow diet during 23 days of the 2^nd^ running period and then HFD was introduced, combined the first two days with chow diet, for 7 days. Chow diet was introduced to evaluate the chow devaluation during exercise. One mouse from inactive mice was also excluded of analysis it run less than 5 km during 1^st^ running period. Statistics were performed without excluded mouse. Despite GEE analysis revealed a main effect of group regarding food intake during the baseline, the pairwise-comparisons did not reveal any differences between groups.

### Study 6: Physiological response of high-runners to HFD during inactivity, measurements and analysis of regional fat

Fourteen 7-week-old male mice were single-housed immediately after arrival and fed with a chow *ad libitum* diet. After 5 days, blocked wheels were provided to 5 mice whereas functional wheels were provided to 8 mice. After 31 days of voluntary wheel running, functional wheels were closed. To avoid increase of food intake during the first days of inactivity impacted on HFD consumption, HFD was introduced together with chow food on day 33. Food preferences were evaluated during day 33-35. Mice were only fed with HFD from day 35 mice. Visceral fat and subcutaneous fat were collected to evaluate the impact of HFD in inactive and sedentary mice.

### Study 7: Molecular analysis of running and long-term inactivity

Fifty-four mice were single-housed immediately after arrival and fed with a chow *ad libitum* diet. Given the mice were used for administration of recombinant proteins (see below), they were split in two cohorts. Sedentary, running and long-inactivity high-runners for 21 days were used for hypothalamic analysis.

### Administration of recombinant proteins

Experiments were performed using 7-week-old single-housed C57BL/6NRj fed with chow standard food. Mice were habituated to injections by receiving once-daily sham injections at least four time before injections of the recombinant proteins before the light off.

### Study 8: Effect of SR59230a on physiological response of exercise

Thirty-eight 7-week-old mice were single housed when arrived and mice were randomized to SR59230a (n=15) or vehicle (n=23) groups based on body weight, body composition and food intake after 2 weeks of arrival. SR59230a (Cat# S8688, Sigma-Aldrich) was resuspended with DMSO at 10mg/ml and diluted with PBS1x. Treated group received a daily injection of 30µg of SR59230a whereas the control mice were administered with PBS1x with the same DMSO concentration (3% DMSO). Three days later and based on food intake and body weight, mice were randomized to sedentary or running groups leading to the following groups: dmso-sedentary mice (n=7), dmso-runners (n=16), SR59230a-sedentary mice (n=7) and SR59230a-runners (n=8). Before the light went off on day 3 and based on food intake and running distance, half a group of dmso-treated running mice were injected with SR59230a (3d-SR59230a-runners; n=8). One mouse from that group died on day 13 and was removed from analysis. Mice were scanned on day 15.

### Study 9: Effect of leptin agonist on exercise-induced food compensation and hypothalamic gene expression

Two cohorts (9.1 and 9.2) of 7-week-old mice were used. In both cohorts, mice were randomized based on body weight, body composition and food intake into sedentary (n=10 in cohort 1 and 2) or running (n=12 in cohort 1 and n=14 in cohort 2). After baseline period, functional wheels were opened, and mice were randomized to leptin agonist or vehicle treatment considering the distance run and food intake on day 1. From day 1 and for 6 days, 5 sedentary and 6 running mice (in the cohort 1) and 5 sedentary and 7 running mice (in the cohort 2) were administered with 30µg of leptin agonist (Cat# 498-OB-05M, R&D Systems) whereas 5 sedentary and 6 running mice (in the cohort 1) and 5 sedentary and 7 running mice (in the cohort 2) were sham-injected with PBS1x. Measurements of body weight, food intake and running distance were taken daily and body composition was measured on day 7 to compare changes produced by leptin agonist in sedentary and running mice. A third cohort (9.3) with 27 mice was used to evaluate hypothalamic gene expression due to running and leptin using the same set-up as beforementioned. On day 3, mice were fasted from 15:00 and injected with leptin or vehicle one hour later. The hypothalami were harvested 90 minutes after injection.

### Study 10: Crossover design of Adiporon administration before and after exercise-induced food compensation

To evaluate the effect of adiponectin agonist (Adiporon; Cat# SML0998, Sigma-Aldrich) before food compensation, mice from cohort 7.2 were administrated with 30 µg of Adiporon in PBS1X or DMSO-PBS1X before day 2 and day 3 of running. Adiporon was diluted with DMSO following the instructions provided by the manufacturer. Mice were injected using a crossover design in which 6 sedentary mice, 7 low-runners and 3 high-runners were injected with DMSO-PBS1X and 6 sedentary mice, 6 low-runners and 4 high-runners were injected with 30 µg of Adiporon before day 2 of running. Next day, those injected with DMSO-PBS1X were injected with 30 µg of Adiporon and vice versa. The same approach was followed for evaluating the effect of Adiporon after food compensation. Mice from cohort 7.1 and 7.2 were administrated with Adiporon and DMSO-PBS1X at 13-16 days of running using a crossover design as above mentioned but resting one day between injections.

### Study 11. αMSH injection

To assess the effect of a-MSH during chronic exercise, sedentary mice and runners from cohort 7.1 were injected with 0.75 µg of analogue of αMSH diluted with PBS1x (Cat# NNC0090-1735, Novo Nordisk) on day 24 and 26 of running period. Mice were injected using a crossover design, being those mice injected with PBS1x injected two days later with a-MSH and vice versa. Only, high-running mice from running group were plotted.

### Tissue and serum collection

Mice were sacrificed by decapitation in the evening near lights off time in *ad libitum* water and food intake. To ensure consistent impact of mouse end-point on study outcomes, mice from each study and group were randomized. However, in case a lot of mice needs to be sacrificed, that was conducted in two days. Blood was collected from trunk and brain was isolated. In some cases, brains were allocated to 4% paraformaldehyde tubes to posterior analysis by immunofluorescence and other cases, fresh hypothalamus was isolated and snap-frozen immediately with liquid nitrogen for posterior analysis. Weights of visceral and subcutaneous fat from the same side were measured and submerged with 4% paraformaldehyde tubes to posterior analysis by immunohistochemistry or snap-frozen immediately with liquid nitrogen to posterior gene expression/protein analysis.

### Serum analysis

Blood was extracted from trunk and centrifugated at 3000g for 10 min. Serum was snap-frozen immediately and stored at −80 °C until used. Mouse adiponectin was analyzed R-PLEX Mouse Adiponectin Antibody Set (Cat# F23YQ-3, MSD) diluting the serum 1000x whereas mouse leptin was assessed by U-PLEX Mouse Leptin Assay, (Cat# K1525ZK-1, MSD) and U-PLEX combo 2 (Cat# K15307K, MSD) diluting the serum 4x from MSD company following the instructions provided by the manufacturer. Mouse Free Fatty Acids were analyzed using a dilution of 4x and Free Fatty Acid Assay Kit (Cat# MAK466, Sigma). Statistical analyses were performed in IBM SPSS Statistics v29.0.1.0. All data are shown as mean ± SEM, and findings with *P ≤ 0.05, **P ≤ 0.01, ***P ≤ 0.001 were considered statistically significant.

### Gene expression analysis

Gene expression profiling in subcutaneous and visceral white adipose tissue of study 4.1 and 4.2 and hypothalamus of cohort 7 and 9.3 was conducted on both sedentary and running mice. Tissues were quickly dissected near lights off time, snap-frozen on liquid nitrogen and stored at -80C until analysis. Total RNA was isolated from tissues with phenol/chloroform extraction method. Briefly, tissue was homogenized with 1 ml of Trizol using Qiagen Tissuelyser Retsch MM300 and spined at 12,000 g for 10 min, 4 °C. Trizol phase was transferred into a new 2ml tubes and 200µl chloroform was added to each sample. After centrifugation, RNA was precipitated using isopropyl alcohol (1:1 reaction), washed with 75% ethanol and dissolved Ultrapure RNase/DNase free water. RNA purity was measured using Nanodrop spectrophotometer (nanodrop one, Thermo Fischer Scientific, USA). RNA was converted into cDNA using cDNA Reverse Transcription Kit (Cat# 4368813, Life Technologies) following the instructions from the manufacturer. Quantitative PCR (qPCR) were performed with PowerUpTM SYBRTM Green Master Mix (Cat# A25780, ThermoFisher Scientific) and 300 nM of forward and reverse primers and using ViiA 7 Real Time PCR system (Applied Biosystems) for amplification. Relative quantification using ΔCt with a fold change normalization at the sedentary group was calculated. *Gadph* was used as housekeeping gene for all tissues. Primers were synthetized by TAG Copenhagen (Table S7). Statistical analyses were performed in IBM SPSS Statistics v29.0.1.0. All data are shown as mean ± SEM, and findings with *P ≤ 0.05, **P ≤ 0.01, ***P ≤ 0.001 were considered statistically significant.

### Transcriptomic analysis by RNA sequencing

RNA from hypothalamus of 7 sedentary mice, 7 high-runner and 8 long-inactivity mice from study 7 were obtained using Trizol-phenol method as previously described and sent to BGI from sequencing. One hypothalamus from high-runner group and 1 from sedentary group were discarded to be sent to sequencing given a technical problem. Stranded RNA-seq libraries were prepared for all high-quality RNA samples using mRNA with poly A tail enrichment. The libraries were then sequenced in paired-end 2x100 nt mode on an DNBSEQ platform. To eliminate batch effects, RNA-seq experiments for samples from the same tissue were performed simultaneously. We generated ∼5 Gb of RNA-seq data for each sample.

First, quality control of the raw sequencing data was performed with SOAPnuke v2.1.5 with default options (*66*). Then, the reads for each of the samples were mapped to the *Mus musculus* genome assembly GRCm39 using STAR v2.7.11b (*67*). The gene counts were retrieved using featureCounts v2.0.6 (*68*), and then built into a matrix. Low-count genes were filtered based on the count-per-million and sample library size (the gene is expressed in at least one replicate). The expression abundance was quantified using cufflinks v2.2.1,(*69*) retrieving the fragments per kilobase of transcript per million mapped reads (FPKM) values. The expression clustering of the replicate samples for each of the sequenced tissues were visualized using PCAs from R package ggplot2 v3.5.1 (*70*). Differential expression analyses were conducted separately for each of the sequenced tissue samples using DESeq2 v1.44.0 (*71*). Volcano plot was used for visualizing each pair-wise comparison of each tissue (sedentary mice vs high-runners, sedentary mice vs inactivity mice and high-runners vs long-inactivity mice) using R v4.4.1 package ggplot2 v3.5.1, dplyr fuction from R package tidyverse v2.0.0 and ggrepel v0.9.6. (*71–73*) Genes with pvalue <= 0.01 and absolute log2FC greater than 1, pvalue <= 0.05 and absolute log2FC greater than 1 and pvalue <= 0.01 were highlighted in red, pink and brown, respectively. Non-significant genes were labelled in grey. We then counted the number of genes with pvalue <= 0.01 from each pair-wise comparison of each tissue and highlighted the overlap of genes between the comparisons using R package VennDiagram v1.7.3 (*74*). The Gene Ontology (GO) functional enrichments of the genes with pvalue <= 0.01 were conducted using enrichGO function of clusterProfiler R package v4.14.3 (*75*). The resulting GO biological process enrichments with adjusted p.value lower than 0.005 were then visualized using R package Rtsne v0.17 and ggplot2 v3.5.1 by labelling the top 10 GO pathways (*76*, *77*). Genes of intersection from each Venn diagram was upload to GOrilla (*78*, *79*).

### Protein analysis of visceral and subcutaneous fat

Approximately 30 mg of tissue was homogenized as previously described,(*80*) with modifications for the extraction of adipose tissue lysate. Briefly, tissue homogenates were prepared using bead mill homogenization (TissueLyser II, Retsch, Haan, Germany), followed by end-over-end rotation at 4°C for 1 hour. To ensure complete removal of adipocyte supernatant, homogenates underwent three consecutive rounds of centrifugation at 18,320 g at 4°C, with subnatant recovery after each round.

Following protein content determination using a Pierce BCA protein assay kit (Thermo Scientific, Rockford, IL, USA), lysates were diluted to equal protein concentrations in ddH2O containing Laemmli sample buffer (62.5 mM Tris-HCl, pH 6.8; 2% SDS; 10% glycerol; 0.1 M dithiothreitol; and bromophenol blue). Samples were heated at 95°C for 5 minutes and subsequently stored at -20°C.

Equal amounts of protein were loaded onto 7-15% SDS-PAGE gels, separated by electrophoresis, and transferred to polyvinylidene fluoride membranes (Immobilion-P, Merck Millipore) using a semi-dry transfer system (Bio-Rad, USA). Membranes were blocked for 30 minutes at room temperature in TBS-T (TBS containing 0.05% Tween-20) supplemented with 3% BSA. Primary antibody incubation was performed overnight at 4°C in either 3% BSA or 2% skimmed milk prepared in TBS-T. The primary antibodies used were: pHSL antibody (Cell signaling, 45804S), HSL antibody (Cell signaling, 4107), AMPKα antibody (Cell signaling, 2532), pAMPKα antibody (Cell signaling, 4185), ATGL antibody (Cell signaling, 2138), TH antibody (ThermoFisher, PA1-4679), UCP1 antibody (Abcam, 10983) and GAPDH (14C10) antibody (Cell signaling, 2118)

The following day, membranes were washed in TBS-T and incubated with horseradish peroxidase-conjugated secondary antibodies for 45 minutes at room temperature. After additional washes in TBS-T, membranes were developed using enhanced chemiluminescence (Immobilion Forte Western HRP substrate, Millipore) and imaged using a ChemiDoc™ MP Imaging System (Bio-Rad, USA). To verify equal protein loading, membranes were washed in water and stained with Coomassie Brilliant Blue.

Images were analyzed using ImageLab software v6.1 (Bio-Rad, USA). Bands corresponding to the protein of interest were quantified using the volume tool and normalized to the average intensity of all bands on the same gel. Statistical analyses were performed in IBM SPSS Statistics v29.0.1.0. All data are shown as mean ± SEM, and findings with *P ≤ 0.05, **P ≤ 0.01, ***P ≤ 0.001 were considered statistically significant.

### Staining and image sample analysis of visceral and subcutaneous fat

Fat tissues were post-fixated using 4% paraformaldehyde and then embedded in paraffin and cut in 2 sections of 2-3µm by 100µm apart by In-lab (Denmark, https://in-lab.dk/). Next, slides were placed inside a heating cabinet at 60°C for at least 1 hour with the aim of melting the paraffin and stained following the Mayer’s Haematoxylin and Eosin Y (H&E) staining protocol. Histological images were acquired using Leica light microscopy DM2000 LED with Leica MC 190 HD camara, and Leica Application Suite X v3.7.6.25997. Small Python v3.10.6 scripts were made to estimate the number and sizes of adipocytes in all sets of images. Python packages used for the image preparation and segmentation were OpenCV, scikit-image, SciPy and Numpy (*81–84*). Python packages used for visualization and data compilation were Matplotlib, Seaborn and Pandas (*70*, *85–87*). All Python scripts and instructions showing how to run them are available at https://github.com/AlexGuBGr/watershed_segmenter. The same functions and pipeline were used for all image sets. However, the images varied in size and resolution across the sets, so each set was analyzed individually to improve the segmentation results. Below, the general preprocessing and segmentation pipeline is described.

#### Preprocessing

Images were read in greyscale and denoised using the denoise_tv_bregman function from Scikit-image. The ridges of the adipocyte borders in the denoised images were detected using the meijering function from Scikit-image. The resulting images were converted to boolean maps (B1), where the detected adipocyte borders were False (background) and everything else was True (foreground). The values used as thresholds between True and False were computed using the threshold_multiotsu function from Scikit-image. To further remove noisy areas in the images, the denoised images were log transformed and processed with the rank.mean function from Scikit-image. The threshold_multiotsu function was then used to compute the additional thresholds distinguishing foreground from background creating additional and separate boolean maps (B2). The two boolean map types (B1 and B2) were elementwise multiplied to get the image structures for the downstream analyses. The combination of B1 and B2 will be referred to as B3. To remove noise, an erosion step followed by a dilation step was performed on B3 images and all connected blobs with a sizes <= 20 were removed.

#### Instance segmentation

To computationally segment the cells into different entities, the watershed algorithm from Skimage was applied. The watershed algorithm treats pixels as topographical features and then “floods” basins (adipocytes) from local minima until all regions meet. If not set, the adipocyte-boundaries will be created where the basins meet.

To create the basins, the Euclidean distances of all pixels to the nearest zero or False-pixels were computed using the distance_transform_edt function from SciPy. The local minima of the individual basins were found on these distance-images using the peak_local_max function from scikit-image, which outputs coordinates of local peaks. We supplied the watershed algorithm with the negative distance-images, B3 images, local minima coordinates and set the watershed_line=True to create the final basin or adipocyte boundaries.

#### Final cleaning step and counting

As a final noise reduction step following the watershed segmentation, all pixels with distances > 0 from all distance-images were collected to create a distance distribution. All pixels with distances to zero pixels higher than the 75th percentile of the distance distribution were marked. The median distances of connected areas (areas not divided by zero or background pixels) within the marked areas were multiplied by their respective sizes (number of pixels) to create another distribution based on median distances and area sizes. This final distribution was log transformed to assume normality, and the lower fence (Q1 -1.5 * IQR) and the upper fence (Q3 + 1.5 * IQR) were computed. All segmented basins that had a log transformed median distance and size relationship below the lower fence or above the upper fence, respectively, were removed. The pixel sizes of the cells in the final instance segmented images were determined and saved as Excel files.

#### Combining image sets and generating plots

To make the sizes of instance segmented adipocytes from the different image sets directly comparable, we normalized the pixel counts such that they became equivalent with imaged based on a 10x magnification. To filter away noise, we removed segmented areas < 200 and > 20000 pixels. Count values of the final datasets and used groups can be found in Table S1. To convert the pixels to 1*10^3 μm^2 we multiplied the pixels counts per cell with 0.042. Boxplots of cell size groups were made in R using the libraries ggplot2 v3.4.3(*77*) and ggsignif v0.6.4(*88*). All statistical analyzes where conducted on the log10 transformed cell size groups using two-sided Mann-Whitney U tests. The set of all p-values per plot were corrected with the Bonferroni method.

### Neuropeptidomics

#### Sample preparation for peptidomic analysis

Hypothalami (∼5 mg) were powdered and lysed in 6 M guanidine hydrochloride buffer (10 mM TCEP, 40 mM CAA, 100 mM Tris, pH 8.5) using Beatbox (PreOmics). Lysates were centrifuged at 4000 g to remove cell debris, and protein aggregation and separation from endogenous neuropeptides was directly induced by adding 80% acetonitrile (CAN; Cat# A955-212, Fisher scientific) to the supernatants. The samples were then centrifuged at 17 000 g for 10 minutes at 4 °C, and supernatants were collected. ACN was removed using a speedvac centrifuge and peptides were then resuspended in a final concentration of 1 % trifluoroacetic acid (TFA; Cat# 8.08260.0101, Sigma-Aldrich) in isopropanol. Peptides were cleaned with 3x styrenedivinylbenzene-reverse phase sulfonate (SDB-RPS; Cat# 66886-U, Sigma-Aldrich) discs using 1% TFA in isopropanol and 0.2% isopropanol in water. Subsequently, peptides were eluted in 60 µl of 1% ammonia and 80% ACN, dried in a speedvac centrifuge and resuspended in 5 µl of 5% ACN and 0.1% TFA. Peptide concentration was determined by a NanoDrop spectrophotometer (Thermo Fisher Scientific) and a total of 150 ng of peptides were loaded on Evotip C18 trap columns (Evosep Biosystems) according to the manufacturer’s instructions.

#### Liquid chromatography mass spectrometry analysis

Peptides were separated using an Aurora (Gen3) 25 cm × 75 μm ID column packed with 1.6 μm C18 beads (IonOpticks) on a Vanquish Neo UHPLC system (Thermo Fisher Scientific). Separation was performed at a constant flow rate of 400 nL/min with a 90-minute stepped gradient: 2–17% solvent B (0.1% formic acid in acetonitrile) over 56 minutes, 17–25% over 21 minutes, and 25–35% over the final 13 minutes. The column temperature was maintained at 50 °C. Eluted peptides were introduced into an Orbitrap Ascend Tribrid mass spectrometer (Thermo) via an EASY-Spray source with a 10 μm fused silica emitter (Evosep) and a FAIMS interface. The spray voltage was set to 1800 V, and the FAIMS compensation voltage (CV) was -45. Data acquisition alternated between full MS scans (120K resolution, 251 ms maximum injection time, AGC target 100%) and 10 data-dependent MS/MS scans (30K resolution, 59 ms maximum injection time, AGC target 400%) using HCD fragmentation. The isolation window was set to 1.4, and the normalized collision energy was 25. To minimize redundant sequencing, selected peptide candidates were excluded from reselection for 60 seconds.

#### Mass spectrometry data processing

Raw MS files were analyzed in FragPipe v22 using MsFragger v4.1 and IonQuant v1.10.27.(*89*, *90*) The search was done using the default parameters in non-specific peptidome workflow against the reviewed mouse fasta reference database UP000000589 (Mus musculus, November 2024).

#### Bioinformatics

Peptidomics data analysis was performed in R v4.3.0 with the following packages: clusterProfiler v4.12.6, dplyr v1.1.4, ggplot2 v3.5.1, factoextra v1.0.7, limma v3.60.6, PhosR v1.14.0, plotly v4.10.4, reshape2 v1.4.4 and tidyverse v2.0.0. Peptide abundances were transformed to log2 scale, samples with low identifications (< 50 %) were removed. Data was filtered for 50 % valid intensity values in at least 1 group (high-runners or sedentary mice). However, more peptide identifications were further removed to ensure that there are at least four samples per group with valid values. In total, we identified 9082 neuropeptides of which 4124 were within >50% of samples. Median normalization was performed. Differential abundance analysis was done using lmFit function by limma package(*91*). The categorization of peptide bioactivity based on different score cut-offs was done using Multipep.(*92*) Correlation analysis of peptide intensities with clinical variables was done using Spearman rank correlations.(*93*) Peptide intensities were imputed using the missForest v1.5 package. For each peptide, Spearman correlations were computed against each clinical variable. Group-specific correlation analyses were conducted by stratifying the dataset based on experimental groups. Gene set enrichment analysis based on biological process, cellular component and molecular function ontologies were done by the ClusterProfiler package,(*75*) retrieving neuropeptide functions from the corresponding parent proteins. For this purpose, the logarithmic fold changes in the limma output file were collapsed using median to create unique identifications at the protein-level. All p-values were adjusted for multiple comparisons using the Benjamini-Hochberg method. Protein interaction network analysis of the shared neuropeptide and peptide hormone proteins was conducted, using STRING functionality v12.0 (*94*).

### Brain immunofluorescence and Image sample analysis

Brains were transferred from the 4 % PFA solution to a 30% sucrose solution at 4C for a minimum of two days until it sinks. Next, they were submerged into freezing 2-methylbutane (Cat# 320404, Sigma-Aldrich) for 30 seconds and stored at -80 ℃ until further processing. Twenty µm brain sections were obtained by cutting coronally the brains and saved at -20C with antifreeze solution (300 mL PB 0,1 M pH 7.2-7.4, 400 mL ethylene glycol (Cat# 102466, Sigma-Aldrich) and 300 mL Glycerol (Cat# G7757, Sigma-Aldrich). Free-flouting immunofluorescence was conducted in the hypothalamic sections. Briefly, sections were blocked with a blocking solution (BS) containing normal donkey (Cat# ab7475, Abcam) serum and Bovine Serum Albumin (Cat# A7906, Sigma-Aldrich) and incubated overnight with primary antibodies at RT. cFOS, ΔFosB, Agrp proteins were detected using Cat# ab190289 (Abcam), 14695S (Cell signaling) and Cat# AF634, respectively. Next day, brain sections were incubated with secondary antibodies for 2 hours at RT in dark room, mounted on slices, covered with permanent mounting medium with DAPI and stored at 4C until analysis. We used secondary antibody donkey anti-Rabbit Alexa 647 (Cat# A-31573, Thermofisher) for detecting cFOS and ΔFosB, and secondary antibody donkey Anti-goat Alexa 555 (Cat# A-31572, Thermofisher) for visualizing Agrp. The detection of cFOS, ΔFosB, Agrp and DAPI was visualized through a fluorescent microscope (EVOS 5000, Invitrogen by Thermo Fischer Scientific) and analyzed manually using CellCounter.jar Fiji ImageJ 1.49 (*95*). To evaluate the chronic and acute neuronal activation of the different hypothalamic areas, study 4.1 and 4.2 were used. Since they were conducted independently, cells per mm2 were calculated and normalized regarding sedentary group. Statistical analyses were performed in IBM SPSS Statistics v29.0.1.0. No statistical methods were applied to predetermine the sample size for experiments. All data are shown as mean ± SEM, and findings with *P ≤ 0.05, **P ≤ 0.01, ***P ≤ 0.001 were considered statistically significant.

## ACKNOWLEDGEMENTS

Anne Jørgensen, Lene Foged and Ida Holm are acknowledged for their technical assistance. This study was supported by a research grant from the Novo Nordisk Foundation (NNF) awarded by C.B (grant ID 0059436) and the Centre for Physical Activity Research (CFAS), which is supported by TrygFonden (grants ID 101390, ID 20045, ID 125132, and ID 177225). P.S is supported by Lundbeck Foundation grant (grant ID R380-2021-1300), N.K. is supported by the BRIDGE – Translational Excellence Programme (Grant ID NNF23SA0087869), supported by the NNF (Grant ID NNF20SA0064340), and the Lundbeck Foundation (Grant ID R436-2023-1225). T.E.J is supported by NNF Ascending Investigator (grant ID 0074481). Proteomics analysis is partially supported by funding from NNF to NNF Center for Basic Metabolic Research (Grant number NNF18CC0034900). Mass spectrometry measurements were performed by the Proteomics Research Infrastructure (PRI) at the University of Copenhagen (UCPH), supported by the NNF (grant agreement number NNF19SA0059305). NNC0090-1735 was provided by Novo Nordisk Compound Sharing.

## AUTHOR CONTRIBUTIONS

Conceptualization, P.S, B.K.P and C.B; Methodology, P.S, B.K.P and C.B; Formal analysis, P.S, J.V, H.H, N.K, K.W.P, A.G.B and J.G; Investigation, P.S, C.B-L, J.V, H.H, N.K, K.W.P, A.G.B, R.S, D.S, and C.B. ;

Visualization, P.S, J.V, H.H, N.K and A.G.B, Writing – original draft, P.S; Writing – review & editing, P.S, C.B-L, J.V, H.H, N.K, K.W.P, A.G.B, R.S, D.S, J.G, T.E.J, A.D, B.K.P and C.B; Supervision, P.S, B.K.P and C.B;

Funding acquisition, P.S, T.E.J, A.D, B.K.P and C.B.

## RESOURCE AVAILABILITY

### Lead Contact

Further information and requests for resources and reagents should be directed to and will be fulfilled by the Lead Contact, Paula Sanchis

## Materials Availability

This study did not generate new unique materials.

## Data and Code Availability

Analysis of subcutaneous and visceral presented in the paper are found Table S1. Scanned western blot images presented in the paper are found Table S2.

The transcriptomic raw RNA-seq data have been deposited at the Sequence Read Archive Database (https://www.ncbi.nlm.nih.gov/sra/) under BioProject accession PRJNA1210847. https://dataview.ncbi.nlm.nih.gov/object/PRJNA1210847?reviewer=c45r9cbjkrm0oqgb74ub3t0ulv

Gene lists and transcriptomic GO data can be found in Table S3 and S4, respectively. Genes used for Venn diagram are named in Table S5.

The mass spectrometry proteomics data have been deposited to the ProteomeXchange Consortium (http://proteomecentral.proteomexchange.org) via the PRIDE partner repository with the data set identifier PXD061648 (https://www.ebi.ac.uk/pride/archive/projects/PXD061648). Neuroropeptidomics processed data can be found in Table S6.

Any additional information required to reanalyze the data reported in this work paper is available from the Lead contact upon request.

## SUPPLEMENTAL INFORMATION TITLES AND LEGENDS

Document S1. Figures S1-S8

Table S1. Data related visceral and subcutaneous cell size. Table S2. Data related to WB analysis

Table S3. Data related genes showed in volcano plots Table S4. Data related GO analysis.

Table S5. Data related Venn diagram Table S6. Data related peptidomics data

Table S7. Data related sequences of primers.

## SUPPLEMENTARY FIGURE LEGEND

**Suppl. Fig. 1. Running drives increased food intake. A**, Simple linear regression between daily food and distance **B-J**) Chronic running in young males fed a control- and calorie-dilution diet. **B**, Schematic. **C**, Lean mass. **D**, Lan mass changes. **E**, Relative fat mass. **F**, Relative fat mass changes. **G**, Lean mass. **H**, Relative lean mass changes **I**, Energy expenditure. **J**, Energy balance. **K-S**) Chronic running in 8- and 26-week-old mice. **L**, Lean mass. **M**, Lean mass changes. **N**, Relative fat mass. **O**, Relative fat mass changes. **P**, Lean mass. **Q**, Relative lean mass. **R**, Energy expenditure. **S**, Energy balance. Two-way repeated-measures ANOVA was used to assess the main effects of running (*) diet (£) or age (#) and its interaction ($) in **C**,**E**,**G**,**I**,**J**,**L**,**N**,**P**,**R**,**S**, but if Mauchly’s and/or Shapiro-Wilk’s test was significant, then repeated-measures GEE was conducted. We applied paired t test to compare data from each mouse group (*) in **D**,**F**,**H**,**M**,**O**,**Q**, and simple linear regression was applied to evaluate the correlations in **A**. Data are mean ± SEM. *P ≤ 0.05, **P ≤ 0.01, ***P ≤ 0.001. **A**, **B** and **K** were created using Biorender.

**Suppl. Fig. 2. Molecular analyses of fat metabolism during short-term *in vivo* experiment. A**, Schematic. **B**, Calories intake. **C**, *Lrrc8Ar* expression. **D**, *Ppary* expression. **E**, *Ki76* expression. **F**, *Fasn* expression. **G**, *Adpr3* expression. **H**, TH protein levels. **I**, Lipe expression. **J**, Proteins levels of pHSL and HSL, and ATGL in visceral fat. **K**, Proteins levels of pHSL and HSL, and ATGL in subcutaneous fat. **L**, Protein levels of UCP1. **M**, Protein levels of pAMPK and AMPK in visceral fat. **N**, Protein levels of pAMPK and AMPK in subcutaneous fat. **O**, Circulating Free fatty acids. **P**, Number of mice used for each group. Independent T-student was used for **A**. One-way ANOVA with simple contrast to reveal differences (*) regarding sedentary group was used for **C**-**P**. In case of data non-normal distribution and/or the heterogeneous variances, GzLM with simple contrast was used. Data are mean ± SEM. *P ≤ 0.05, **P ≤ 0.01, ***P ≤ 0.001.

**Suppl. Fig. 3. Molecular analysis of hypothalamus as well as leptin and Adiponectin from short-term and chronic running. A**, Acute neuronal activation in ARC. **B**, Acute neuronal activation in other hypothalamic areas. **C**, Acute neuronal activation in VMH subareas. **D**, Chronic neuronal activation in VMH subareas. **E**, Agrp staining in the hypothalamic areas. **F,** Circulating leptin and adiponectin in chronic running mice. **G,** Gene expression in the visceral fat. **H**, Gene expression in the subcutaneous fat. I, Protein levels in visceral fat. **J**, Protein levels in subcutaneous fat. **K**, Association between circulating leptin levels and relative visceral and subcutaneous fat weight at ZT0 and ZT12. **L**, Association between circulating leptin levels and circulating free fatty acids. One-way ANOVA with simple contrast was used to reveal differences (*) regarding sedentary group in **A**-**E**,**G**-**J**. One-way ANOVA with Bonferroni post hoc multiple comparisons was assessed in **F**. However, if data did not follow a normal distribution and/or the variances were not homogeneous, GzLM with simple contrast for running groups regarding sedentary group or sequential Bonferroni post hoc multiple comparisons was used instead respectively. Simple linear regression was applied in **K,L.** Data are mean ± SEM. *P ≤ 0.05, **P ≤ 0.01, ***P ≤ 0.001.

**Suppl. Fig. 4. Further phenotypic analysis related to Fig. 4. A**-**D**) Daily leptin injection study**. A**, Schematic. **B**, Relative fat mass. **C**, Relative lean mass. **D**, Hypothalamic gene expression **E**-**G**) Crossover design with Adiporon during chronic running. **E**, Schematic. **F**, Total distance. **G**, Total intake. Two-way ANOVA was used to assess the main effects of running (*), drug () and its interaction ($) in **B**-**D**. If data did not follow a normal distribution and/or the variances were not homogeneous, GzLM was used. Paired T-test was applied for assessing the effect of each group (*) in **F**,**G**. Data are mean ± SEM *P ≤ 0.05, **P ≤ 0.01, ***P ≤ 0.001. **A** and **E** were created using Biorender.

**Suppl. Fig. 5. Further phenotypic analysis related to Fig. 5. A-J**) Effect of age during inactivity. **A**, Schematic. **B**, Body weight. **C**, Lean mass. **D**, Lean mass changes. **E**, Relative fat mass. **F**, Associations between phenotypic characteristics. **G**, Association of intake and lean mass. **H**, Energy expenditure. **I**, Energy expenditure changes. **J**, Energy balance. **K**-**T**) Effect of long-term inactivity. **K**, Schematic. **L**, Lean mass. **M**, Lean mass changes. **N**, Relative fat mass. **O**, HFD exposure and withdrawal. **P**, Food preferences. **Q**. Overconsumption of HFD on 1^st^ day of exposure. **R**, Overconsumption of HFD period. **S**, HFD-related chow devaluation on 1^st^ day of chow exposure. **T**, HFD-related chow devaluation on last experimental day (*left*) and association chow devaluation and running (*right*). **U**, Food preferences during inactivity. One-way repeated-measures ANOVA was conducted to evaluate the main effect of running (*) in **L**,**N** and two-way repeated-measures ANOVA was conducted to evaluate the main effect of running (*), age (#) and its interaction ($) in **B**,**C**,**E**,**H**,**J**. In case of non-normal distribution and/or that sample’s sphericity was significant, GEE with sequential Bonferroni post hoc multiple comparisons was used. One-way ANOVA was used to evaluate the main effect of running (*) in **O-T** and two-way ANOVA was used to evaluate the main effect of running (*), age (#) and its interaction ($) in **D**,**I**. However, in case of non-normal distribution and/or heterogenous variances, GzLM with sequential Bonferroni post hoc multiple comparisons was used. Independent T-student was used for **U** and simple linear regression was used for **G**, **I** and **T**. Data are mean ± SEM. *P ≤ 0.05 **P ≤ 0.01, ***P ≤ 0.001. **A**, **O** and **Z** was created using Biorender.

**Suppl. Fig. 6. Further analysis related to Fig. 6. A**, Volcano plots of hypothalamic transcriptome. **B**, Correlations between peptides’ log (intensities) and clinical data. **C**, Gene set enrichment analysis based on molecular function. Icon from BioRender were used in **A**.

