## Supplementary figures and images for "The metabolic and molecular mechanisms underlying running-induced energy compensation"

### Suppl figures

Suppl. Figure 1

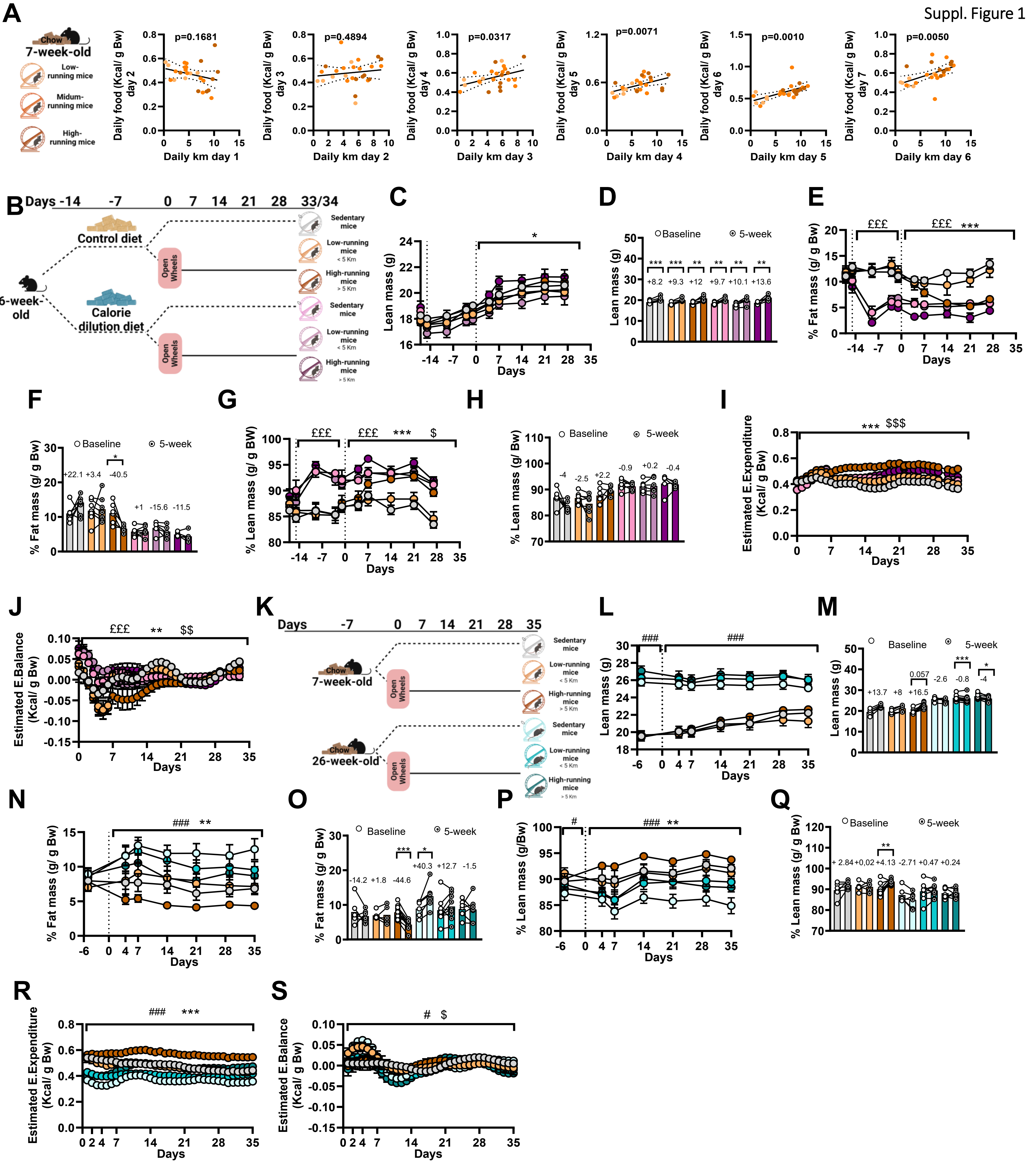

Suppl. Figure 2

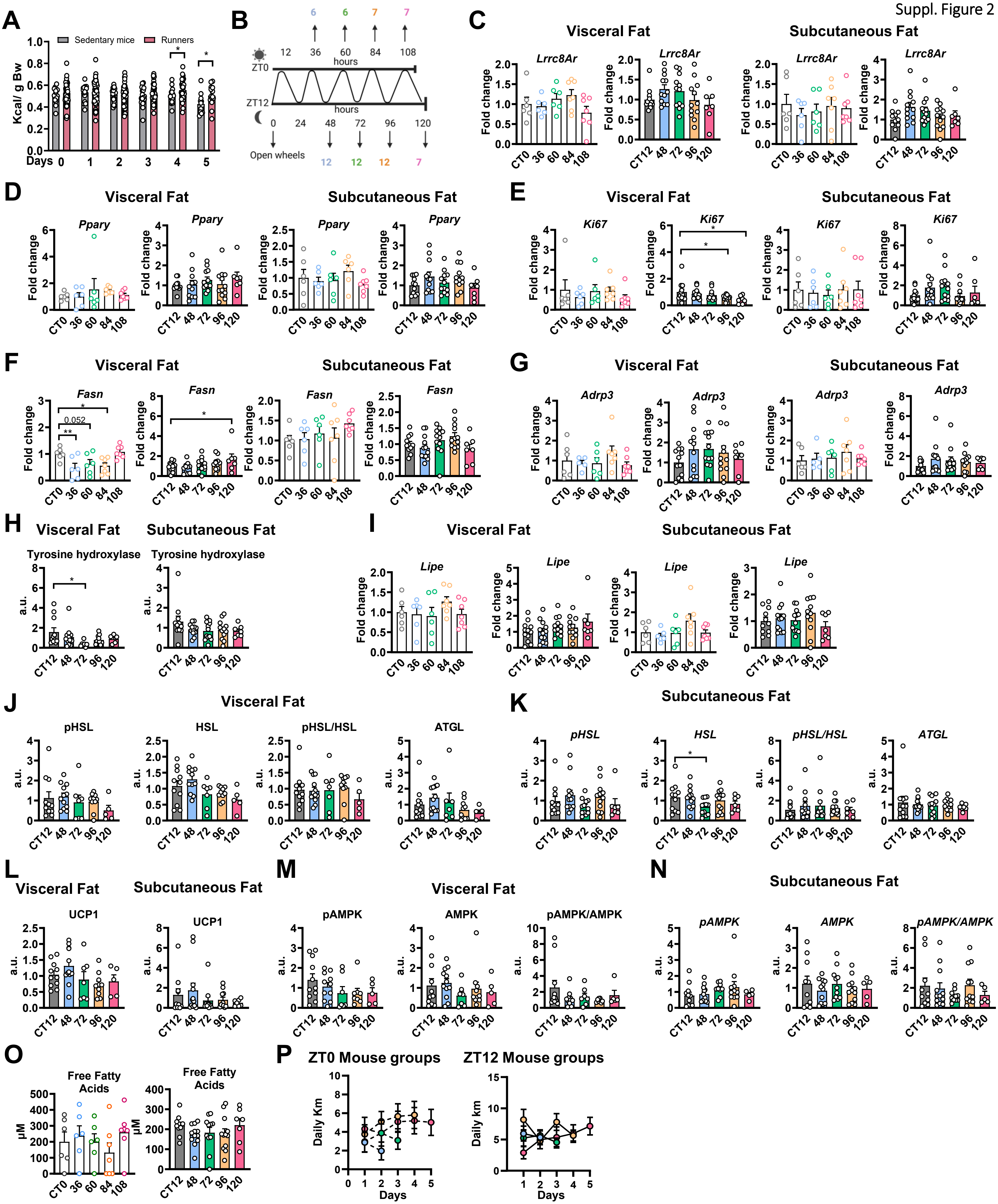

Suppl. Figure 3

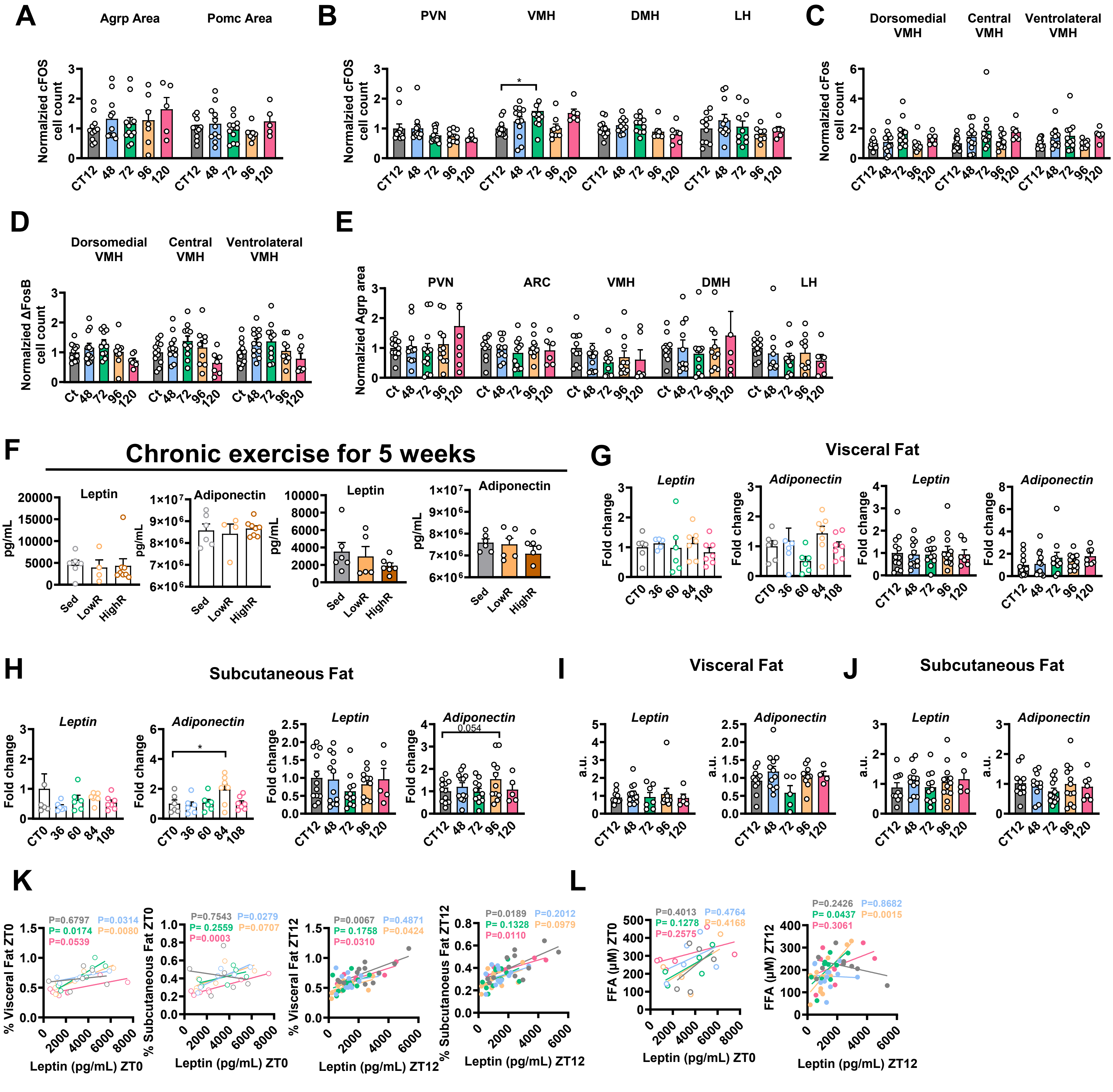

Suppl. Figure 4

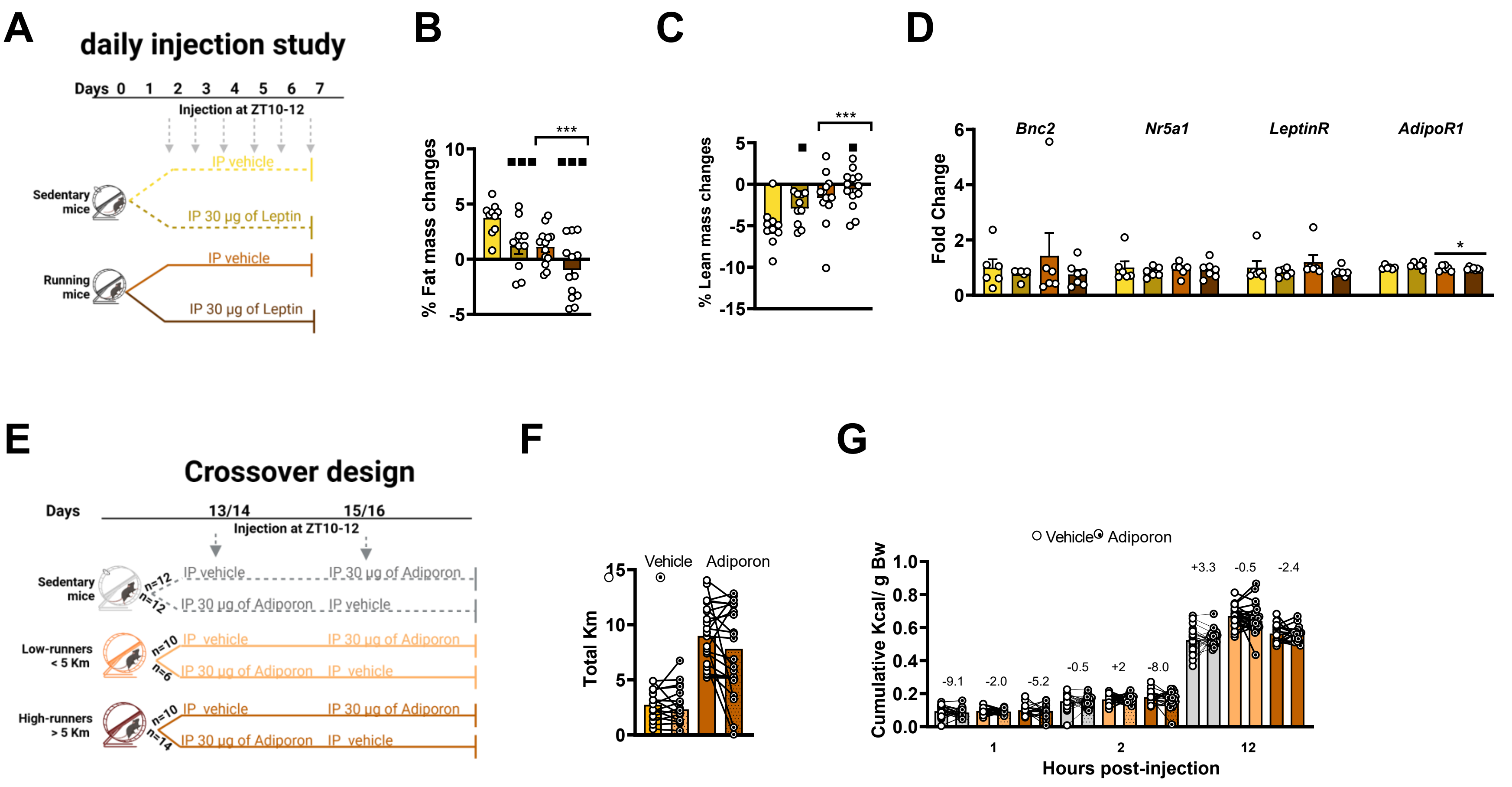

Suppl. Figure 5

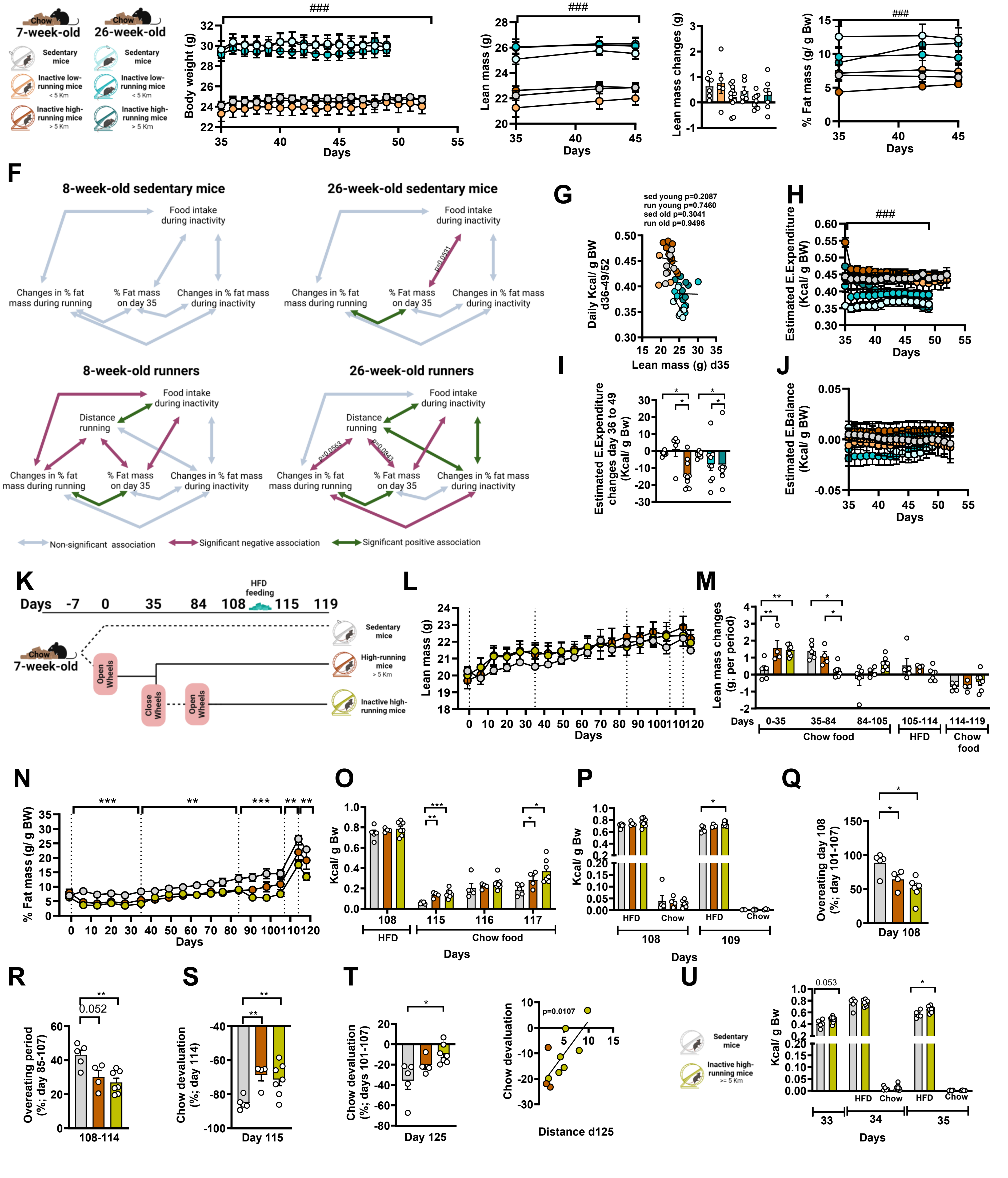

Suppl. Figure 6

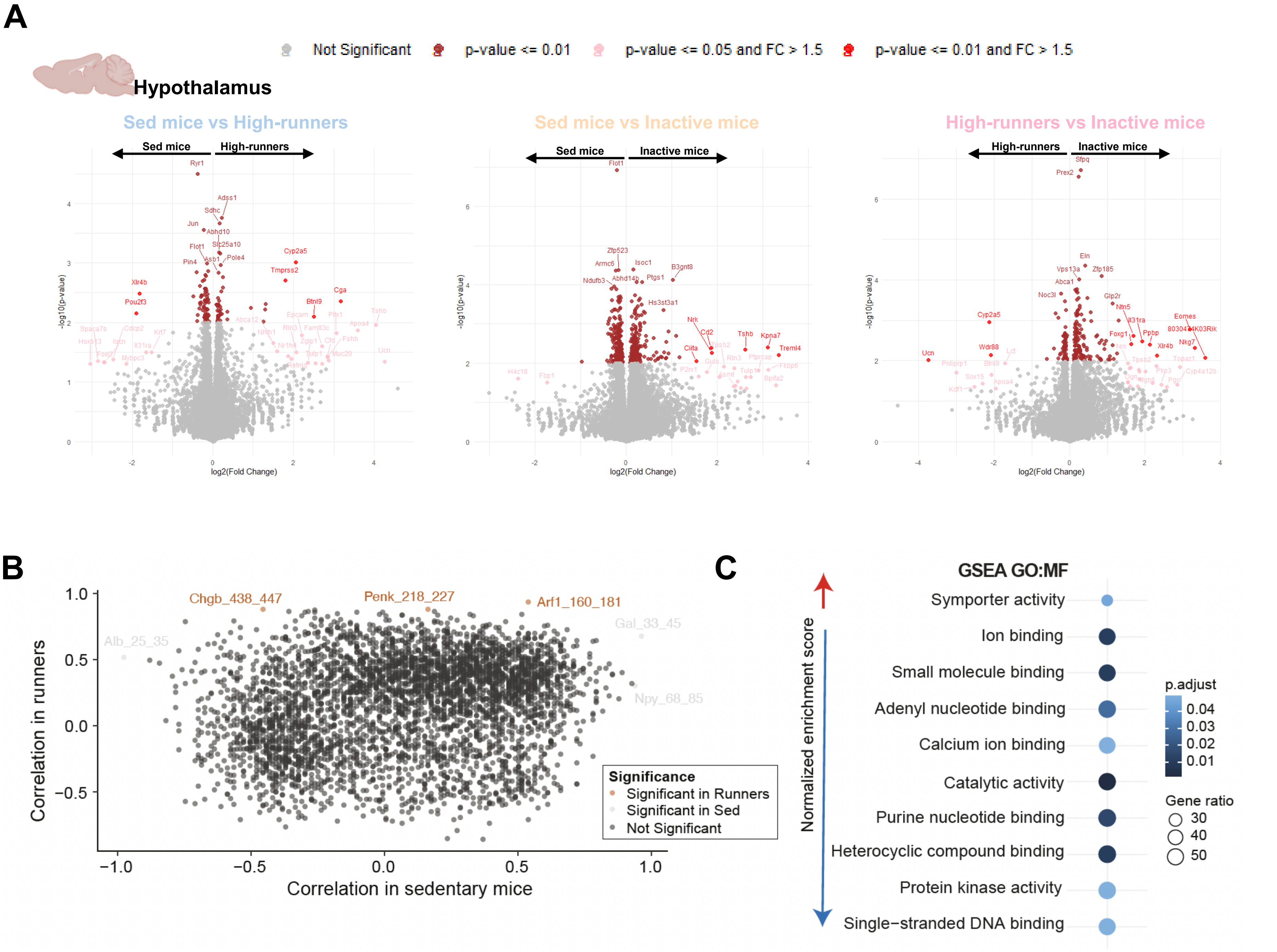
